# PAK1, PAK1Δ15, and PAK2: similarities, differences and mutual interactions

**DOI:** 10.1101/580928

**Authors:** D. Grebeňová, A. Holoubek, P. Röselová, A. Obr, B. Brodská, K. Kuželová

## Abstract

P21-activated kinases (PAK) are key effectors of the small GTPases Rac1 and Cdc42, as well as of Src family kinases. In particular, PAK1 has several well-documented roles, both kinase-dependent and kinase-independent, in cancer-related processes, such as cell proliferation, adhesion, and migration. However, PAK1 properties and functions have not been attributed to individual PAK1 isoforms: besides the full-length kinase (PAK1-full), a splicing variant lacking the exon 15 (PAK1Δ15) is annotated in protein databases. In addition, it is not clear if PAK1 and PAK2 are functionally overlapping. Using fluorescently tagged forms of human PAK1-full, PAK1Δ15, and PAK2, we analyzed their intracellular localization and mutual interactions. Effects of PAK inhibition or depletion on cell-surface adhesion were monitored by real-time microimpedance measurement. We show that PAK1Δ15 is in many aspects similar to PAK2, rather than to PAK1-full. Both PAK1Δ15 and PAK2, but not PAK1-full, were enriched in focal adhesions, indicating that the C-terminus might be important for PAK intracellular localization. Using immunoprecipitation, we documented direct interactions among the studied PAK group I members: PAK1 and PAK2 form homodimers, but all possible heterodimers were also detected. Our results indicate that PAK1 and PAK2 have distinct roles in cell adhesion and mutually affect their function. PAK1-full is required for formation of membrane protrusions, whereas PAK2 is involved in focal adhesion assembly. We have also noted that PAK inhibition was associated with a large reduction of the cell glycolytic rate. Altogether, our data suggest a complex interplay among different PAK group I members, which have largely non-redundant functions.

## Introduction

PAK (p21-activated kinases) are a group of serine-threonine kinases originally identified as downstream effectors of p21 proteins, specifically of the Ras-related GTPases Rac1 and Cdc42 (1, 2). The initial phase of PAK research focused on their role as small GTPase effectors in the context of dynamic remodelling of the cytoskeleton and of cell adhesion structures (3). Later on, PAK were found to be involved in many cancer-related processes in different tumor types (4, 5). In parallel, the discovery of PAK1 nuclear localization (6) prompted the analysis of PAK functions in the cell nucleus.

The human PAK family is divided into the group I (PAK1 to PAK3) and group II (PAK4 to PAK6). In general, these kinases regulate the cytoskeleton dynamics, intracellular signaling, and gene expression (7). The current knowledge about PAK group I is derived mostly from adherent cell models, where PAK activity usually correlates with increased cell motility. In this area, the research was mainly focused to PAK1, although some important differences between PAK1 and PAK2 have been reported (8). In addition, PAK are known regulators of a wide range of cellular processes, including the dynamics of actin structures and microtubules, cell division, apoptosis, and adhesion to the extracellular matrix.

Despite considerable sequence homology within PAK group I, the individual members appear to have distinct function in cell physiology. Whereas PAK2 is more or less ubiquitiously expressed, PAK1 expression is more restricted in adulthood, and PAK3 is expressed only in the brain (9). PAK2 and PAK4 are essential during embryonic development, since knockouts are embryonic lethal, at least in mice (9).

Group I PAK share some domains that are not present in the group II members (10, 11). In particular, the autoinhibitory domain (AID) is important for regulation of the kinase activity of the group I family members. The regulatory mechanism was described on the basis of PAK1 crystal structure (12), and is supposed to be valid also for PAK2 and PAK3. PAK1 forms homodimers in a *trans* arrangement, the AID of one molecule interacting with the kinase domain of its partner molecule (13, 14). In this closed conformation, the kinase activity is very low. Binding of a small GTPase, like Rac1 or Cdc42, to the p21-binding domain (PBD) of PAK1 triggers conformation changes in the kinase domain, leading to dimer dissociation and to subsequent changes of conformation associated with increasing kinase activity (15).

Phosphorylation at Ser223 during this process was reported to be required for full PAK1 activation (16). Autophosphorylation at PAK1 Ser144, or at the equivalent site for the other PAK, stabilizes the open conformation and sustains high kinase activity. Apart from p21 proteins, PAK1 activity can be regulated by PxxP motif of SH3 domains (17) or through phosphorylation by Akt (18), JAK2 (19, 20), CDK1/cyclin B1 (21), PDK1 (22), or other kinases, which often regulate binding of phospholipids or scaffold molecules like GRB2 or NCK1.

Whereas PAK1 has been the predominant PAK group I member in studies focusing on the cell adhesion and migration, PAK2 was mainly studied in association with its role in the apoptosis. Upon PAK2 cleavage at a consensus caspase-3 site, the N-terminal fragment (28 kDa), containing the AID, dissociates from the C-terminal part (34 kDa), presumably inducing constitutive kinase activity (23). Interestingly, *in vitro* studies suggest that the cleaved PAK2 molecules could remain in dimers (24). Cells expressing a dominant-negative PAK2 were still able to undergo the apoptosis, but morphological changes, like membrane blebbing and formation of apoptotic bodies, were inhibited (23, 25). PAK2 cleavage induced by cellular stress occurs in a caspase-dependent manner (26–28), and contributes to apoptosis in various cell types. On the other hand, the full-length PAK2 was reported to have anti-apoptotic effects (29–31).

Human cancer is usually not associated with PAK mutation, but rather with a dysregulated PAK expression (7), especially with PAK1 and PAK4 overexpression. Both PAK1 and PAK4 genes are found on chromosomal regions that are frequently amplified in cancer (32). PAK1 is the most studied and upregulated in cancers arising from PAK1-expressing tissues, such as brain, pancreas, colon, or ovary (7). PAK activity has been linked to uncontrolled cell proliferation, altered cellular signaling, increased metastasis formation, and regulation of the immune system. PAK overexpression was also associated with resistance to several drugs like paclitaxel, doxorubicin, cisplatin, and 5-fluorouracil (7, 33).

Given their role in tumor-related processes, PAK were proposed as possible targets in anti-cancer treatment (34–38). However, with regard to specific functions of different family members, it will be necessary to search for more specific PAK inhibitors or to inhibit specific downstream effectors. This will require further studies of signaling pathways related to the individual PAK family members. Functional differences between PAK1 and PAK2 in relation to cell adhesion have been described in a human breast carcinoma cell line, using small interfering RNAs (8). Although both PAK1 and PAK2 contributed to increased cell invasiveness, their roles were mediated by distinct signaling mechanisms. In addition, possible diversity is not limited to different family member genes: PAK1 has at least two confirmed splicing isoforms, denoted as PAK1A and PAK1B in the Swiss-Prot database, and little is known about their respective functions. Compared to the full length PAK1B (the transcript variant 1 according to RefSeq database, PubMed), the variant PAK1A (transcript variant 2) lacks the exon 15. In a melanoma model, overexpression of the shorter isoform (denoted as PAK1Δ15) did not trigger MAPK signaling and had no effect on the cell proliferation rate, in the opposition to the full length form (39). Furthermore, altered ratio between PAK1 and PAK1Δ15 in melanoma patients was associated with more aggressive disease and with worse prognosis.

In the present work, we compared the properties of the full-length PAK1, PAK1Δ15 and PAK2, focusing on their respective roles in the processes involved in the cell adhesion.

## Results

### Antibody characterization

The expression level of PAK in HEK293T and HeLa cells was analyzed using a set of antibodies (Table 1). Whereas PAK2 was detected in a single dominant band at about 60 kDa, multiple bands were observed between 60 and 70 kDa using anti-PAK1 detection. Fig. 1 shows a direct comparison of signals obtained using selected antibodies. At least three bands attributable to PAK1 were identified on large gels, with different relative intensities for different antibodies. In general, the bands detected by phospho-specific antibodies were at slightly higher position compared to the total protein bands. The antibody recognizing the autophosphorylation site pSer144/141 on PAK1/PAK2, respectively, showed at least three PAK1 bands, which will be denoted as pPAK1-0, pPAK1-1 and pPAK1-2 in this study (Fig. 1). PAK3 was not detected in any of the cell lines studied.

**Table 1.** PAK antibodies

| Cat. No | Producer | Target | Immunogen | Type |
| --- | --- | --- | --- | --- |
| #2602 | Cell Signaling | PAK1 | N-terminal peptide | rabbit polyclonal |
| ab40852 | Abcam | PAK1 | N-terminal peptide | rabbit monoclonal |
| ab223849 | Abcam | PAK1 | peptide within<br>AA 200-300 | rabbit monoclonal |
| ab131522 | Abcam | PAK1 | region around<br>AA 210-214<br>(P-V-T-P-T) | rabbit polyclonal |
| ab76293 | Abcam | PAK2 | N-terminal peptide<br>AA 1-100 | rabbit monoclonal |
| ab40795 | Abcam | PAK group I<br>pSer144/141 | sequence around the<br>autophosphorylation<br>site in PAK1 (pSer144) +<br>PAK2 (pSer141) + PAK3<br>(pSer139) | rabbit monoclonal |
| ab75599 | Abcam | PAK1 pT212 | sequence around<br>phosphorylated Thr212,<br>not present in PAK2 | rabbit polyclonal |

**Fig. 1:**
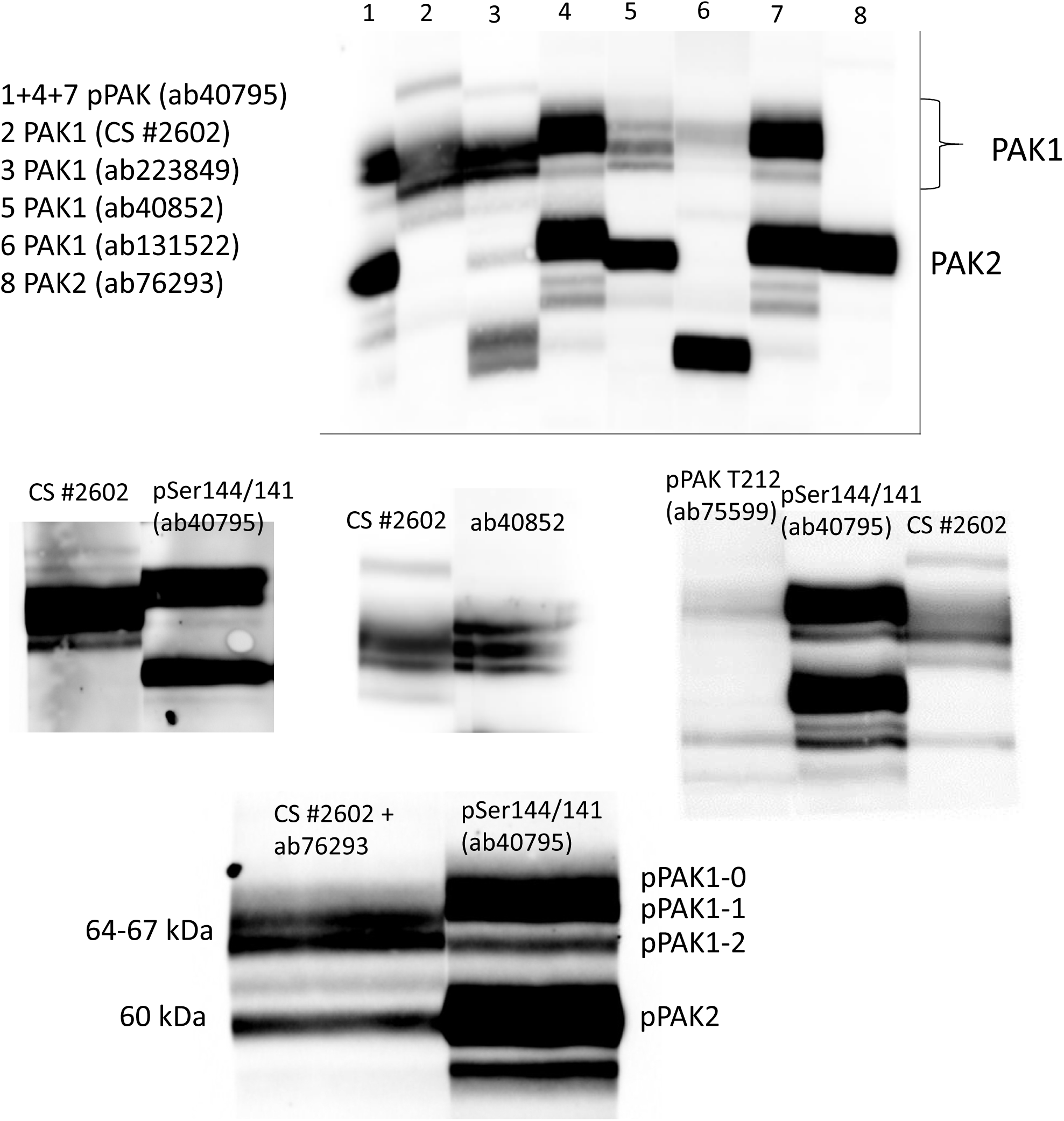
Comparison of signals from different PAK antibodies. HEK293T cell lysate was resolved on a large gel, the proteins were transfered to a nitrocellulose membrane. The membrane was vertically cut, the individual parts were incubated with the indicated primary antibody, then with the corresponding secondary antibody. The membrane was reassembled, covered with the chemiluminiscence substrate and the signal was recorded from all the parts at once. Detailed description of antibodies used is given in Table 1.

The specificity of the antibodies was further checked using siRNA-mediated PAK1/PAK2 silencing. As it is shown in Fig. 2, all bands between 64 and 70 kDa were specific for PAK1, with the exception of those detected by ab131522, which gave no specific signal from the naturally occurring PAK. In addition, several strong unspecific signals were also identified for other antibodies, at lower molecular weight (MW).

**Fig. 2:**
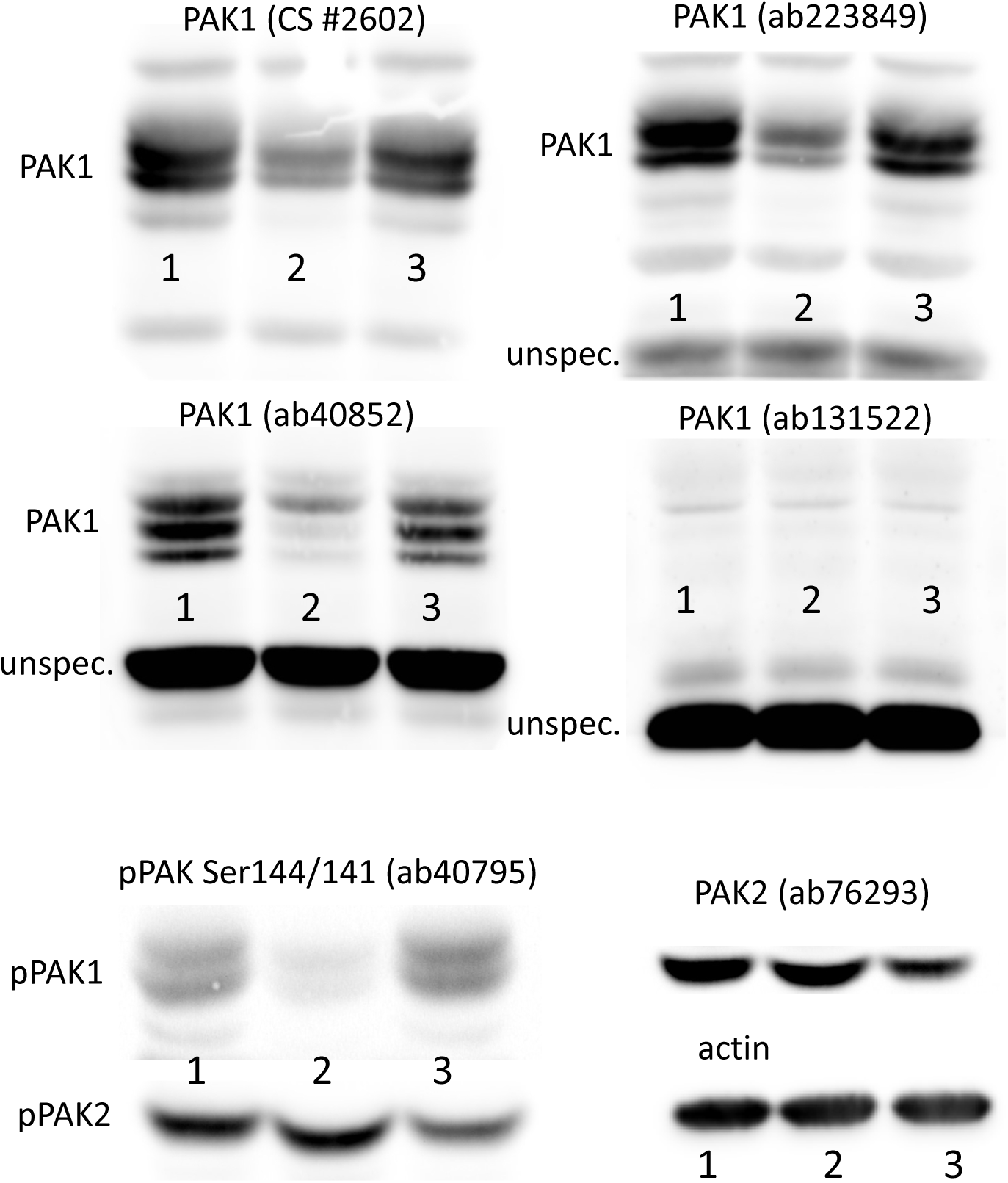
Effect of siRNA PAK1/PAK2 on western-blot band intensity (antibody specificity control). HEK293T cells were transfected with siRNA targeting PAK1 or PAK2 and incubated for 48 h. The cell lysates were resolved on 18×18 cm gels and PAK expression was detected using different antibodies as indicated. Lanes: 1 – untransfected control, 2 – siRNA PAK1, 3 – siRNA PAK2.

### PAK1 variants

To further explore the nature of the observed PAK1 bands, we have constructed plasmids for exogenous expression of two PAK1 splicing variants (39): the full-length isoform (PAK1-full) and a shorter variant lacking the exon 15 (PAK1Δ15). The sequence comparison of these two isoforms, along with PAK2 sequence, is given in the Supplementary Information (Fig. S1). Some important sequence differences and the percentage of sequence homology for the individual domains are illustrated in Fig. 3. Due to a frameshift, the C-terminal part of PAK1Δ15 is different from that of PAK1-full. On the other hand, there is substantial similarity between the C-termini of PAK1Δ15 and PAK2. All PAK group I have an autoinhibitory domain (marked as “AID” in Fig. 3), which includes a dimerization sequence (“dimer”), and the kinase domain (“kinase”). The phosphorylatable Threonine 212 (T212) is present in PAK1 variants, but not in PAK2. On the other hand, PAK2 is unique in having a cleavage site for caspases (“casp3”), followed by a myristoylation site (“myr”) (40).

**Fig. 3:**
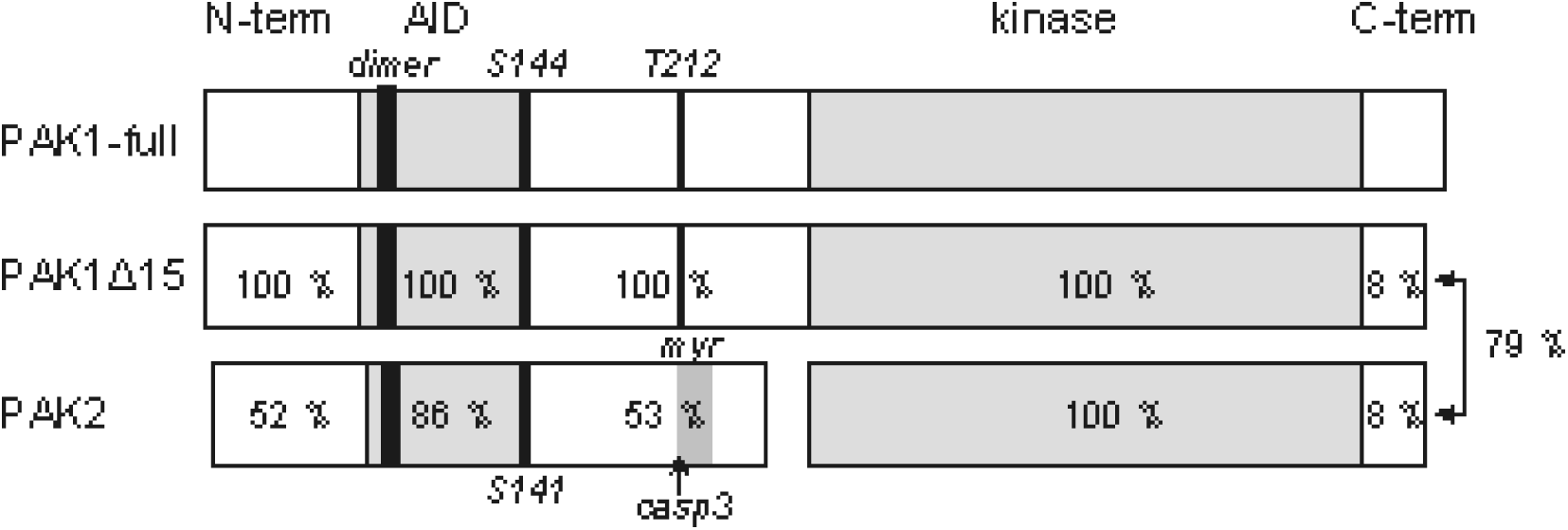
Schematic illustration of PAK group I structure. The numbers within the individual PAK1Δ15 and PAK2 domains indicate the sequence homology with the corresponding domains of the full-length PAK1. The dimerization sequence (dimer, AA 79-86) is identical for all three isoforms. The figure depicts the position of some important elements: the autoinhibitory domain (AID), the kinase domain (kinase), the autophosphorylation site S144/S141, the phosphorylation site T212, the cleavage site for caspase-3 (casp3) and the myristoylation region (myr). The C-terminal domains of PAK1Δ15 and PAK2 are similar to each other (79% homology), but different from that of PAK1-full.

Fig. 4 shows the position of new bands formed in cells transfected with plasmids coding for PAK1-full or PAK1Δ15. As all the used PAK1 antibodies target the N-terminal half of the protein, they should recognize both these variants equally. Indeed, the pattern of the exogenous bands was similar and the main difference was a shift towards lower MW due to the shorter C-terminus of PAK1Δ15. The exogenous kinases appear little phosphorylated at Ser144, as no increase in the signal intensity after transfection was observed using the corresponding phospho-specific antibody. Interestingly, the exogenous proteins were also detected by the antibody ab131522, which did not recognize the endogenous PAK1 forms in HEK293T cells. On the other hand, only the higher bands from the exogenous products were apparent using the ab40852 antibody.

**Fig. 4:**
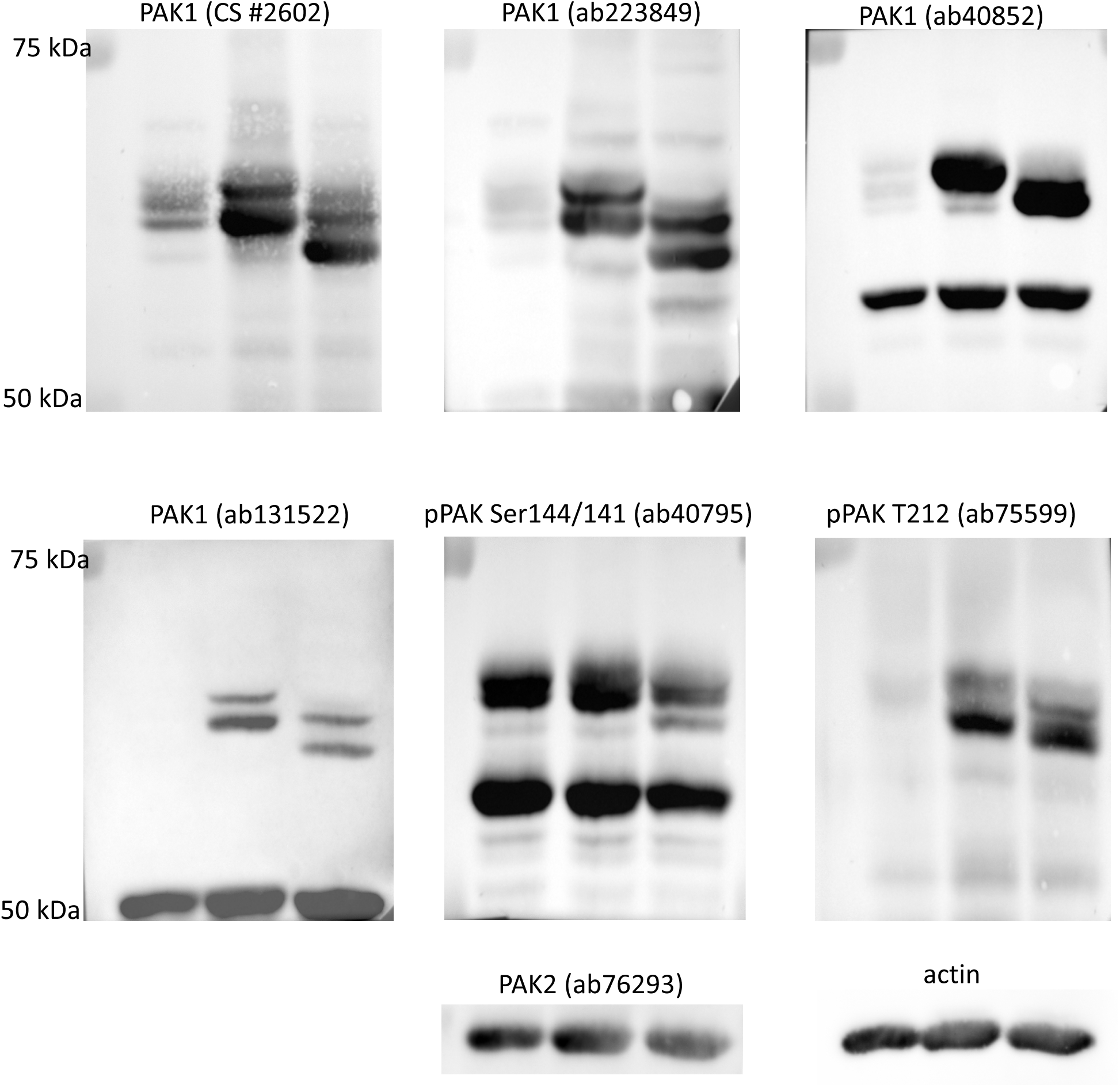
Band positions for the exogenous PAK1 forms. HEK293T cells were transfected with plasmids coding for PAK1-full or PAK1Δ15, incubated for 48 h and lysed. The proteins were resolved on 18×18 cm gels and blotted to nitrocellulose membranes, which were incubated with different antibodies as indicated. Lanes: 1 – untransfected cells, 2 – exogenous PAK1-full, 3 – exogenous PAK1Δ15

Both plasmids induced formation of at least two distinct bands. Previously, it was reported that PAK1 activation is associated with a change of its electrophoretic mobility, the activated form having higher apparent MW on polyacrylamide gels. The shift is generally supposed to be due to a hyperphosphorylation occurring during kinase activation. To verify this explanation, we analyzed the effect of alkaline phosphatase (AP) treatment of cell lysates on the band pattern (Fig. 5). The efficiency of AP treatment was confirmed by phospho-specific antibodies: signals from both pSer144/141 and pT212 were largely reduced in all AP-treated samples. However, the position of PAK1 bands was not substantially altered by the treatment. Instead, Fig. 5 suggests that the affinity of some of the PAK1 antibodies is affected by phosphorylation. Although a slight shift was usually detectable for the exogenous PAK1 forms, the pattern was not substantially modified by AP treatment. The nature of different products formed from the transfected plasmids is thus not clear. Double bands were also obtained for fluorescently labeled PAK1 variants (see later).

**Fig. 5:**
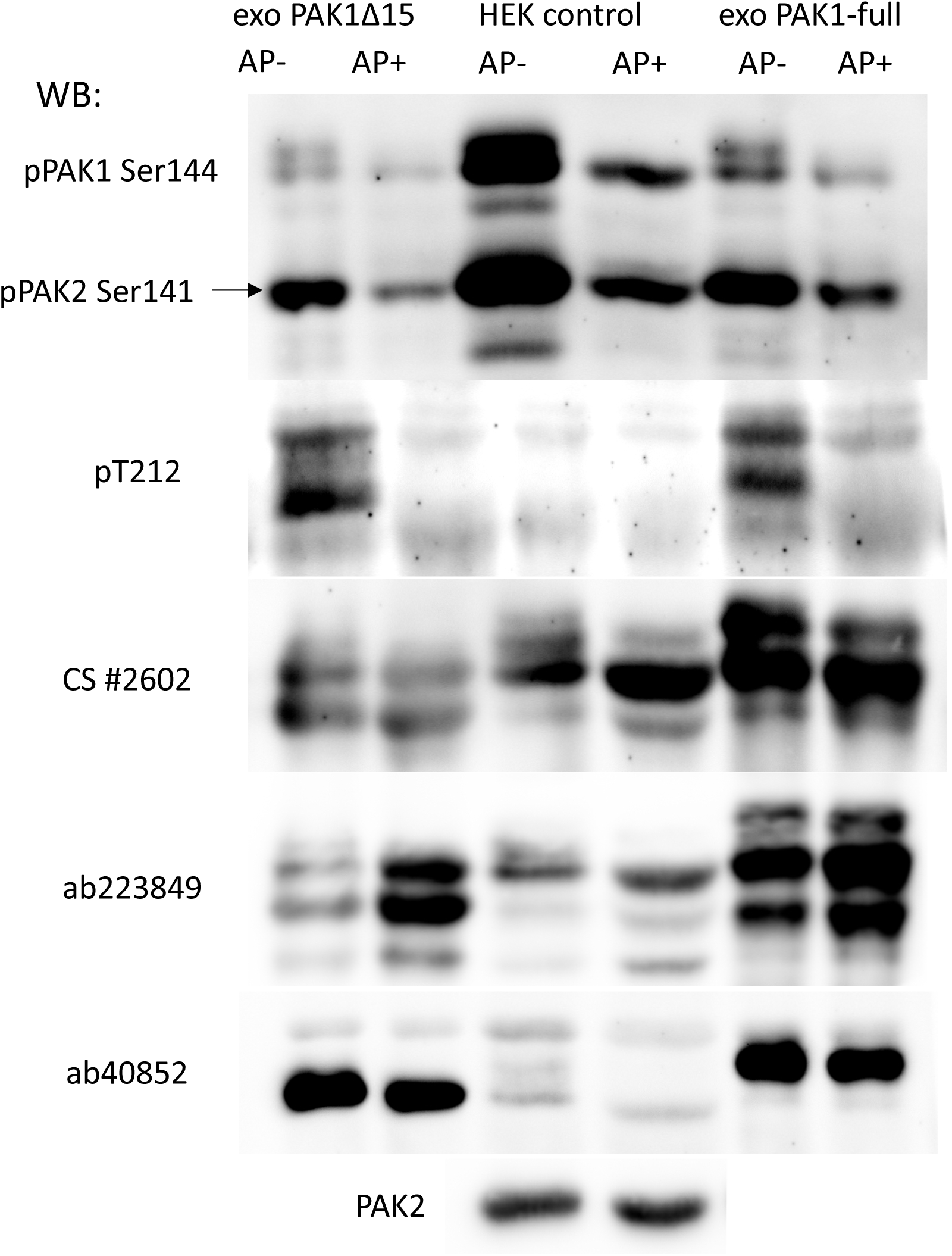
Effect of alkaline phosphatase treatment on western-blot bands of PAK. Lysates from HEK293T cells, control or transfected with plasmids coding for PAK1-full or PAK1Δ15, were incubated overnight with alkaline phosphatase (AP). The proteins were resolved on 18×18 cm gels and blotted to nitrocellulose membranes, which were incubated with different antibodies as indicated. The total protein load was five times lower in the lanes containing transfected cells than in those containing the control cells.

### PAK dimerization

PAK1 is known to form homodimers in a *trans* arrangement, where the kinase domain of one molecule is bound to AID of the other one. The kinase activity is low in this conformation and binding of a small GTPase to PAK1 regulatory domain is required for dimer dissociation and kinase activation. PAK2 homodimers are supposed to be formed as well, on the basis of sequence similarity and of indirect experimental confirmation (24). However, no data are available as to possible heterodimer formation involving different members of PAK group I family. We thus constructed plasmids allowing for expression of eGFP- or mCherry-tagged PAK variants and analyzed their mutual interaction by co-immunoprecipitation. The exogenous proteins with their interaction partners were pulled down from HEK293T cell lysates using GFP/RFP Nano-Traps, and the presence of the co-precipitated forms was assessed through the complementary label (anti-RFP/GFP antibody, respectively). The representative western-blots from these experiments are shown in Fig. 6. The panel A documents the expected homodimer formation for PAK1-full, but also a strong interaction between PAK1-full and PAK2. Surprisingly, the formation of PAK1/PAK2 heterodimer was associated with PAK2 cleavage: although the GFP-labeled PAK1 is only slightly larger than the GFP-labeled PAK2, a marked difference between the position of the co-precipitated PAK1 and that of the co-precipitated PAK2 was found systematically (cf lane 2 versus lane 1 in the blot IP:RFP, WB:GFP). The truncated form of PAK2-GFP was also detectable in the cell lysates, exclusively in samples with PAK1+PAK2 co-expression, but its amount was low compared to that of the intact PAK2-GFP (cf lane 2 with the other lanes in the blot input, WB: GFP). In the RFP-immunoprecipitates, the fragment represented the dominant PAK2-GFP form: see Fig. S2 for direct comparison of the band position in the lysate and in the precipitate. Identical results were obtained using the inverse labeling, i.e. mCherry-labeled PAK2 and eGFP-labeled PAK1. Again, the interacting PAK2 was cleaved whereas PAK1-full was intact (data not shown). The length of the truncated PAK2 was reminiscent of the apoptotic fragment (p34), which is formed upon caspase-mediated PAK2 cleavage. As eGFP was added from the C-terminal side, the resulting fragment would have about 66 kDa.

**Fig. 6:**
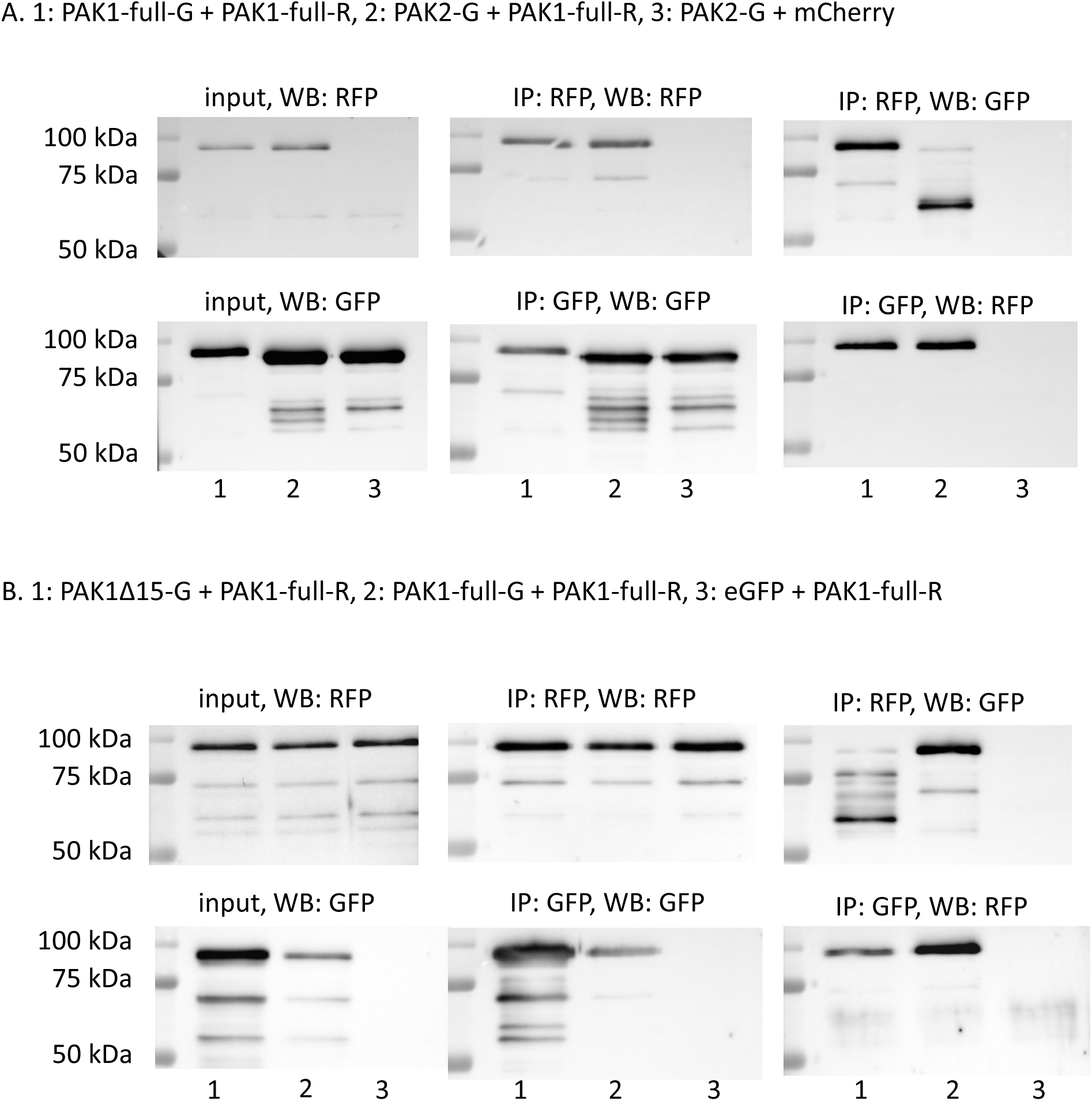
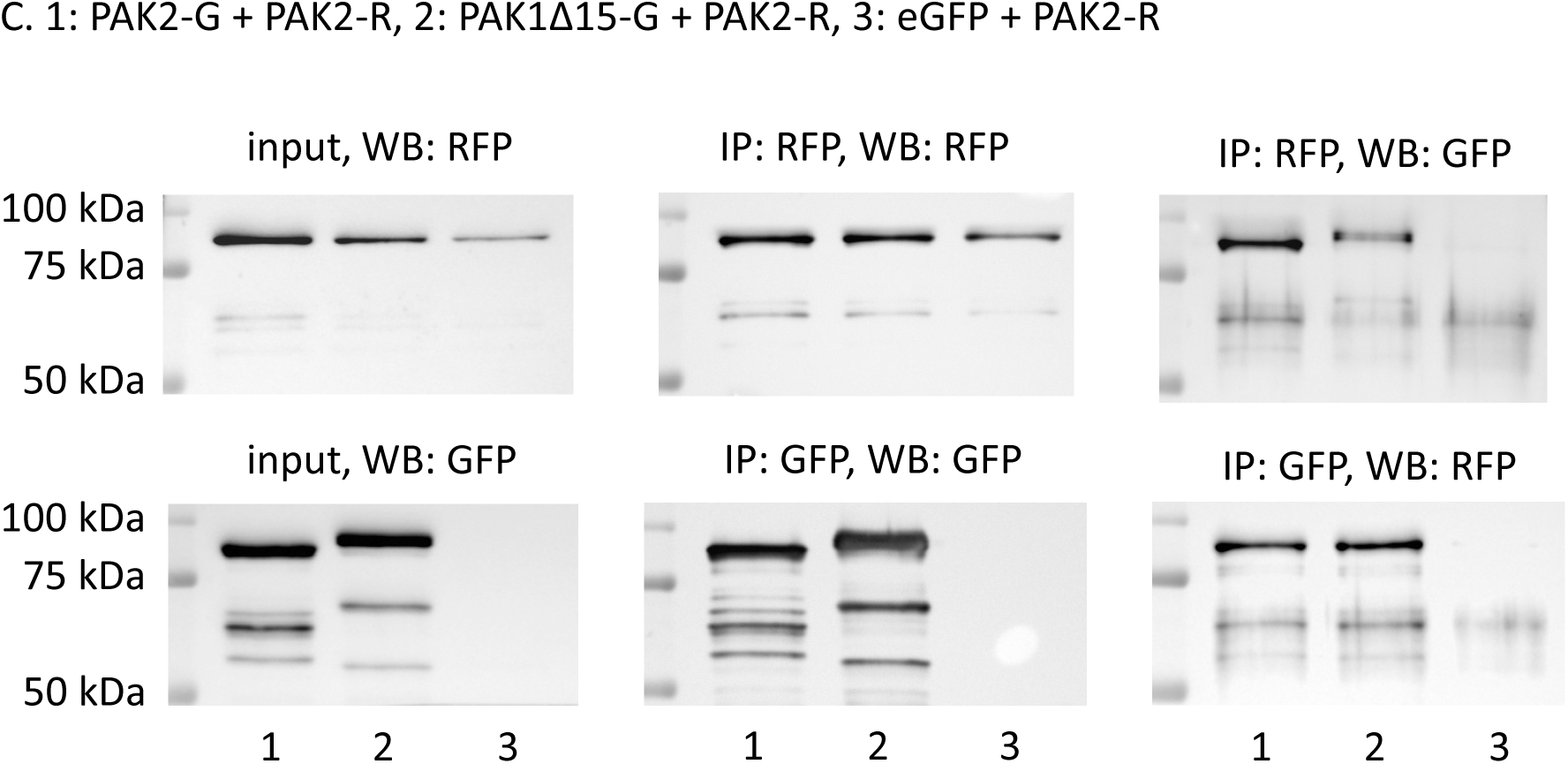
Results of co-immunoprecipitation experiments. HEK293T cells were cotransfected with combinations of labeled PAK isoforms (G-green variant, R-red variant) and empty eGFP/mCherry plasmids. Proteins were precipitated through GFP or RFP beads (indicated as IP:GFP or IP:RFP). The membranes were probed with anti-GFP or anti-RFP antibodies (indicated as WB:GFP or WB:RFP). The input (lysate) is also shown in the left column. MW markers are shown on the left of each blot.

However, cell pretreatment with the caspase inhibitor Q-VD-OPh did not attenuate the cleavage (Figure S3). Using the same approach, we found that PAK1-full interacts with PAK1Δ15, the latter kinase being cleaved in association with the complex formation (Fig. 6, panel B). On the other hand, no cleavage was observed for PAK2 interaction with PAK1Δ15 (Fig. 6, panel C). We have also directly proved that PAK2 forms homodimers (Fig. 6, panel C, lane 1). The summary of the detected complexes is provided in Table 2.

**Table 2:** Summary of direct interactions among PAK group I proteins

|  | PAK1 full-length | PAK1Δ15 | PAK2 |
| --- | --- | --- | --- |
| PAK1 full-length | homodimers detected by co-IP | interaction detected by co-IP, PAK1Δ15 cleavage | interaction detected by co-IP, PAK2 cleavage |
| PAK1Δ15 |  | homodimers indicated by native electrophoresis | interaction detected by co-IP, no cleavage |
| PAK2 |  |  | homodimers detected by co-IP |

The search for possible mixed endogenous/exogenous complexes was complicated by the presence of shorter products formed from the plasmids. Nevertheless, the comparison of images obtained from anti-GFP and anti-PAK1 antibodies suggests that the endogenous PAK1 binds to PAK1-GFP (Fig. S4).

Complex formation was further analyzed using native electrophoresis. Lysates from cells transfected with PAK1-full-eGFP, PAK1Δ15-eGFP or PAK2-eGFP were prepared and resolved under non-denaturating conditions, transferred to a membrane and tested using anti-GFP antibody (Figure 7). The position of PAK1Δ15 was only slightly higher compared to PAK2, in agreement with slightly larger MW. On the other hand, PAK1-full was not detectable in the native gels, probably because it was bound in multimolecular complexes, that are too large to enter the gel. The presence of PAK1-full in the cell lysates was confirmed under denaturing conditions (SDS electrophoresis), where the proteins migrate as monomers. So far, the observed properties of PAK1Δ15 were often close to those of PAK2. However, both PAK1-full and PAK1Δ15, but not PAK2, were always detected in double/multiple bands.

**Fig. 7:**
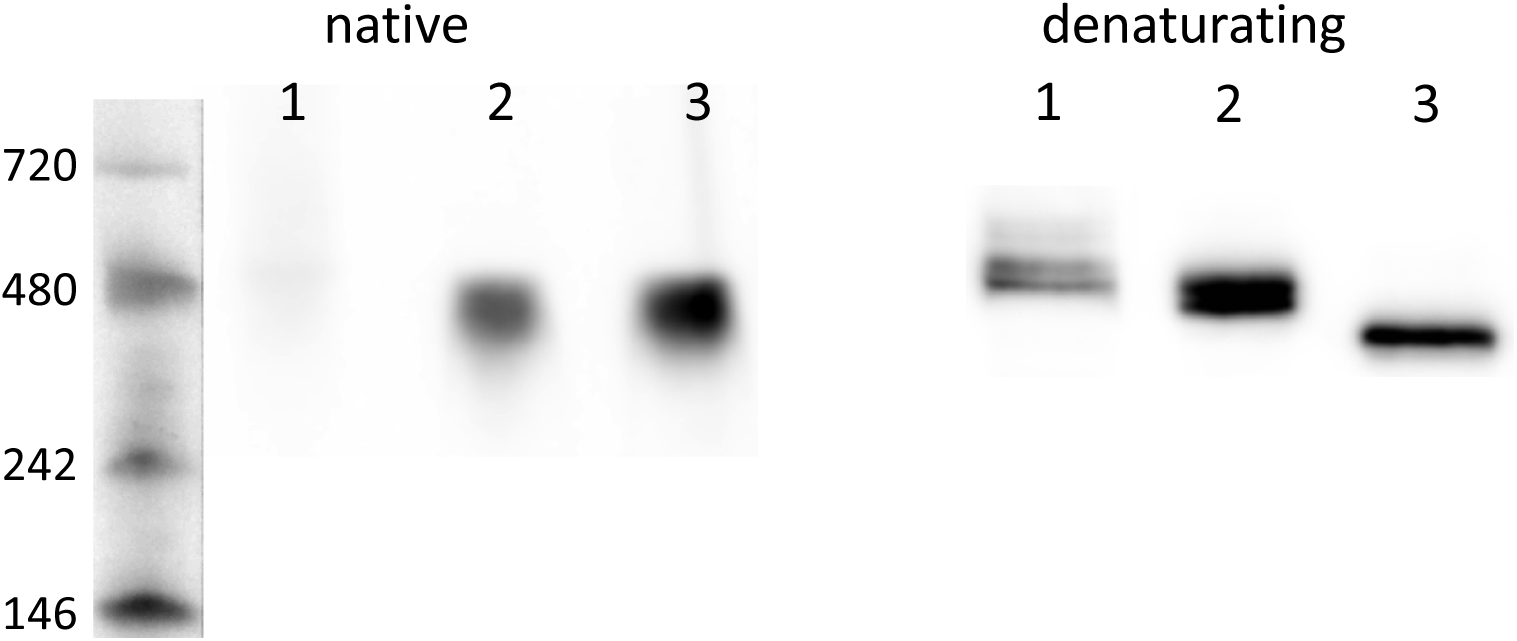
Analysis of PAK complex formation using native electrophoresis. HEK293T cell lysate from cells transfected with plasmids encoding PAK1-full-GFP, PAK1Δ15-GFP or PAK2-GFP were resolved on electrophoretic gels under native or denaturating conditions. MW markers (kDa) are shown on the left for the native gel. Lanes: 1 – PAK1-full, 2 – PAK1Δ15, 3 – PAK2.

### Intracellular localization

The fluorescently labeled forms of PAK1-full, PAK1Δ15 and PAK2 were further used to study the intracellular localization of the individual PAK group I members. The analysis was usually performed 24 to 48 h after HeLa cell transfection with the corresponding plasmids. For PAK1-full, we observed mainly a diffuse cytoplasmic staining, the signal being more intense in the proximity of the plasma membrane. No accumulation of the fluorescence signal in adhesion points was noted. In contrast, both PAK1Δ15 and PAK2 were clearly enriched in focal adhesions, which were visualized either by paxillin/vinculin staining or using the interference reflection microscopy (IRM) (Figure 8A). When the cells were co-transfected with fluorescently labeled PAK1-full and PAK2/PAK1Δ15, the signal from PAK1-full was found beyond the adhesion points labeled with PAK2/PAK1Δ15 (Fig. 8B). This corresponds to the known function of PAK1 in regulating dynamic actin structures inducing membrane protrusions.

**Fig. 8:**
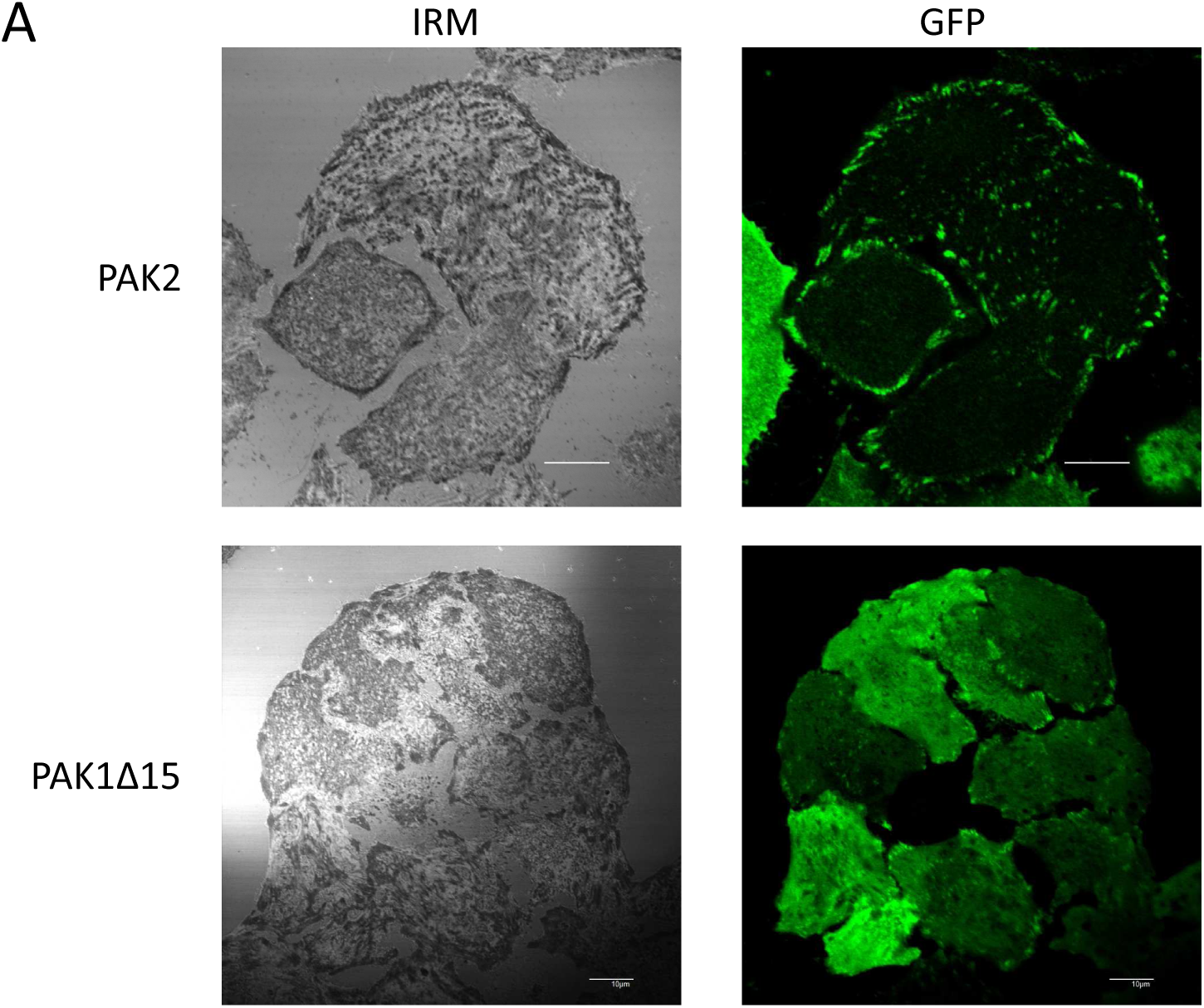
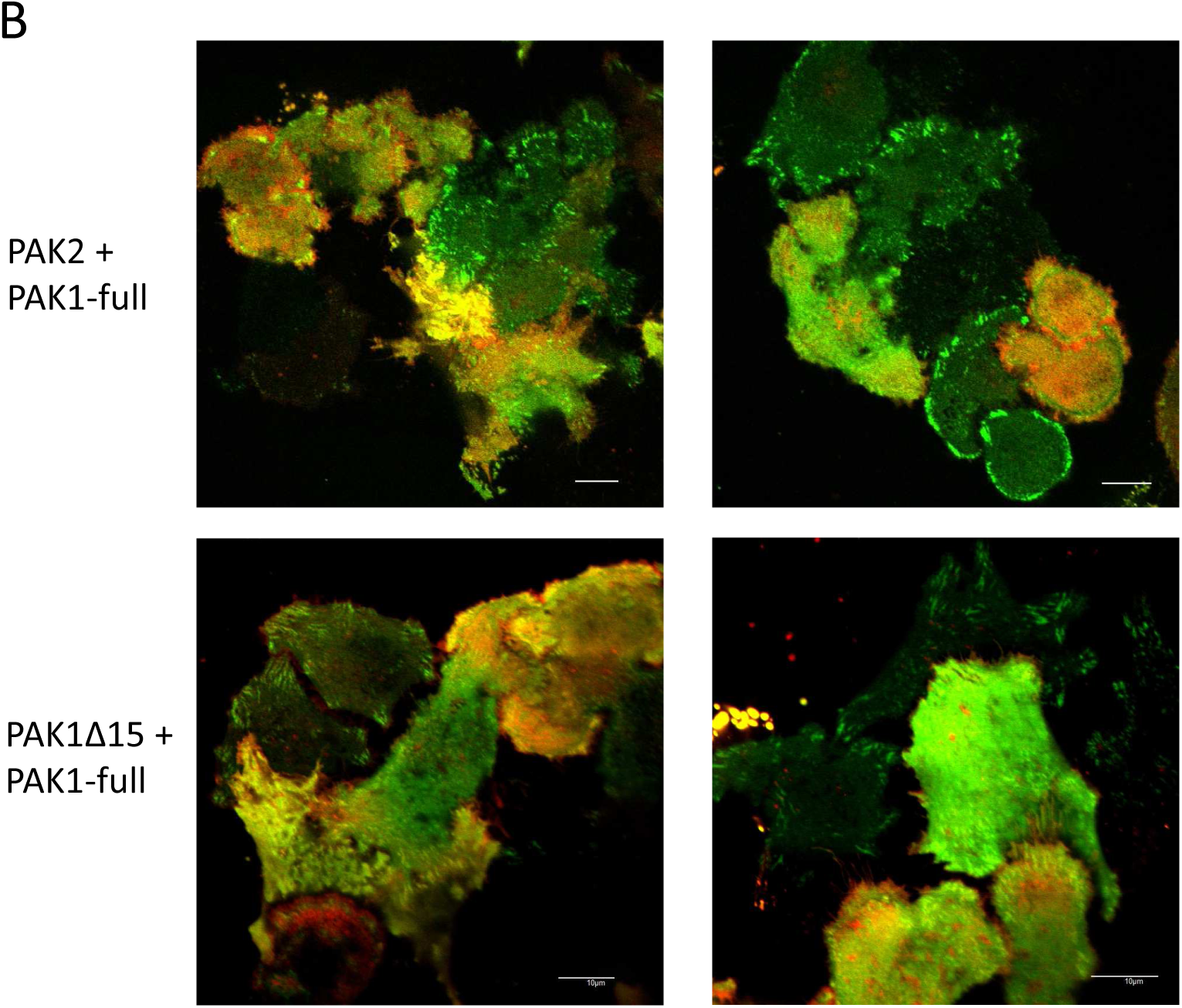
Analysis of PAK intracellular localization. HeLa cells were transfected with plasmids encoding fluorescently labeled variants of PAK1-full, PAK1Δ15 or PAK2, and the intracellular localization of the exogenous proteins was analyzed using the confocal microscope FluoView 1000 (Olympus). A: Localization of PAK1Δ15-eGFP and PAK2-eGFP in cell-surface adhesion points, which were visualized by the interference reflection microscopy (IRM). Scale bars: 10 µm. B: Analysis of the intracellular localization of PAK1Δ15/PAK2 (green) and PAK1 full-length (red) in co-transfected cells. The scale bars correspond to 10 µm.

Endogenous PAK were visualized by immunofluorescence. The only antibodies suitable for this application were ab76293, which is specific for PAK2, and the phospho-specific (pSer144/141) antibody detecting the autophosphorylated (kinase-active) forms of both PAK1 and PAK2. As PAK1 expression level in HeLa cells was found to be very low (Fig. 9), the signal of the phospho-specific antibody is mainly attributable to PAK2 in HeLa cells. Immunofluorescence staining confirmed the presence of PAK2 in adhesion structures of HeLa cells (Fig. 10A). Moreover, we observed an intense staining of microtubule organizing centers (MTOC) in mitotic cells (Fig. 10B). In both HeLa and HEK293T cells, MTOCs were clearly labeled by pSer144/141 antibody, but not by PAK2 antibody. PAK1 binding to microtubules and to MTOC has already been observed previously and was related to T212 phosphorylation (21). We did not detect any clear MTOC staining using the exogenous fluorescently labeled proteins. However, in HeLa cell line, mitotic cells were not present in the cell subpopulation expressing these proteins.

**Fig. 9:**
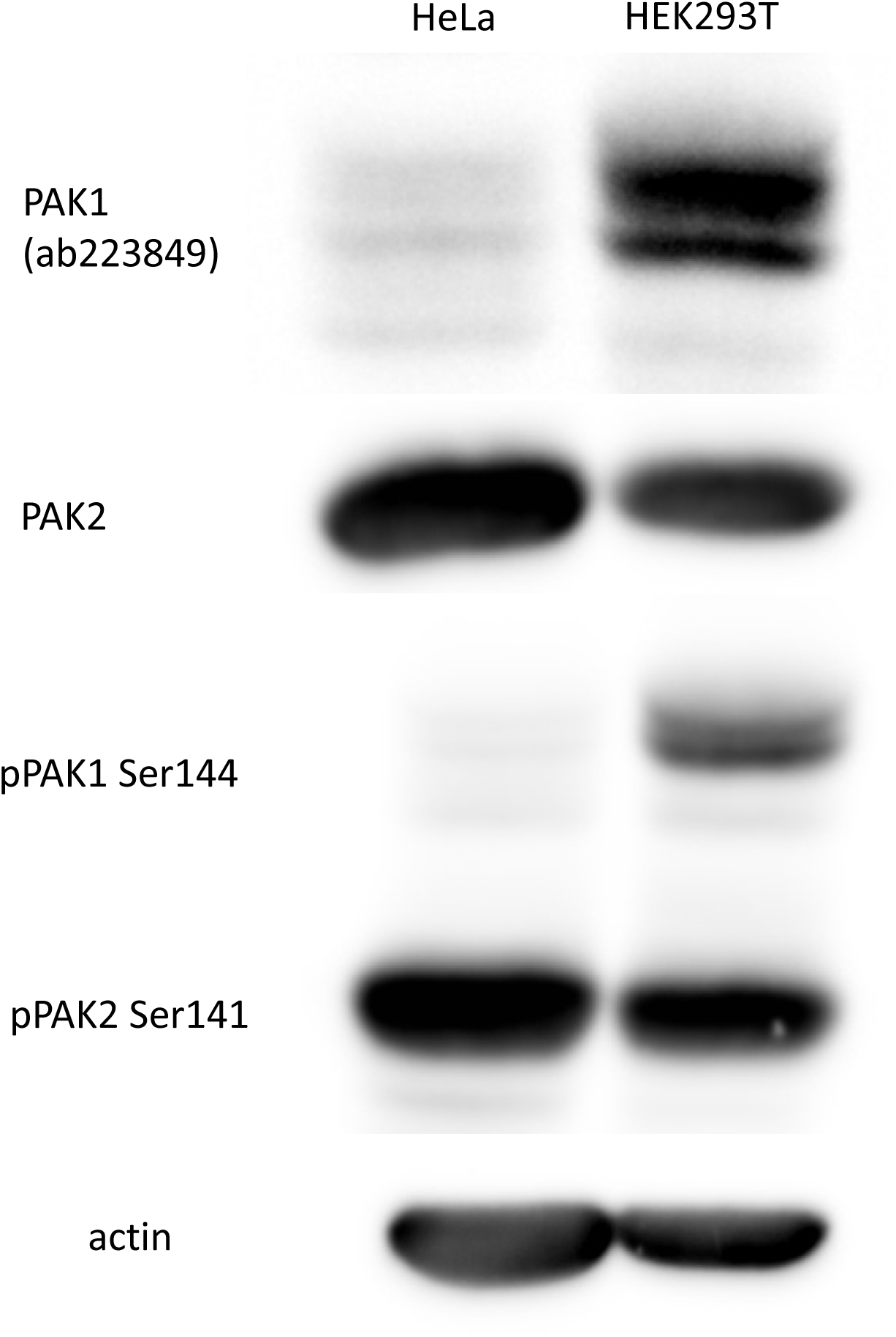
Comparison of PAK1 and PAK2 expression level in HeLa and HEK293T cells. Lysates from HEK293T and HeLa cells were resolved in 18×18 cm gels, the proteins were transferred to a membrane and visualized using the indicated antibodies.

**Fig. 10:**
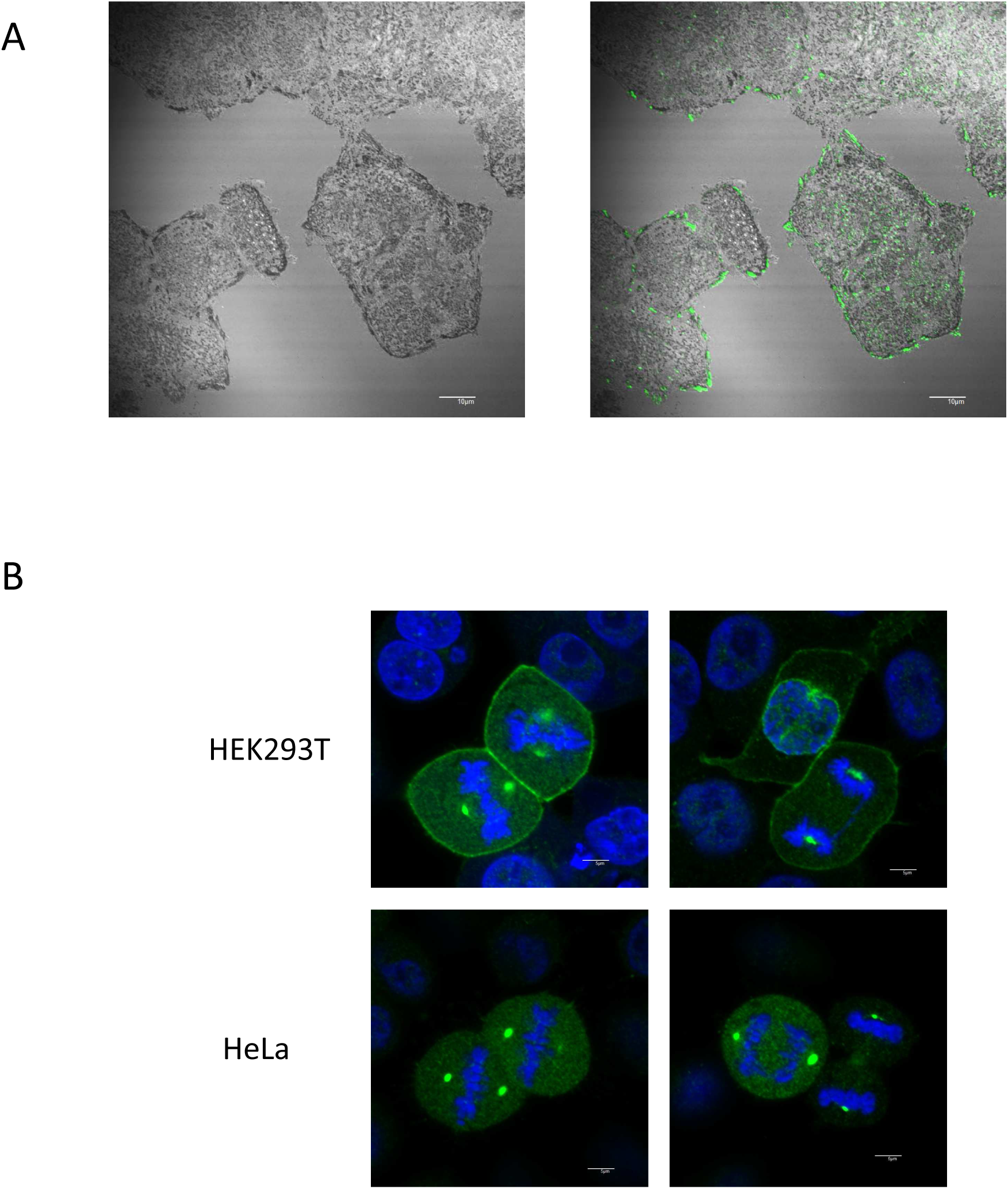
Analysis of endogenous PAK localization by immunofluorescence staining. A. Left: HeLa cell-surface adhesion points visualized by interference reflection microscopy (IRM). Right: PAK2 signal (green) is superposed on IRM signal from the same visual field. B. Microtubule organizing centers stained by the phospho-specific (Ser144/141) antibody to PAK1/PAK2 in mitotic cells. Green – pPAK1+pPAK2, blue – nuclei. Scale bars: 10 µm.

### Effect of small molecule inhibitors

To analyze the impact of an acute inhibition of PAK, we employed the compound IPA-3, which binds covalently to the regulatory domain of PAK group I proteins. The efficiency of IPA-3 depends on the cell density (41) and we thus adhered to identical conditions in all experiments. Moreover, as IPA-3 causes oxidative stress, it is recommended to use a control compound, PIR3.5, which also induces oxidative stress, but does not alter PAK activity. We have verified that the toxicity of both inhibitors was similar over the used concentration range (Fig. S5).

PAK1 is a known downstream effector of kinases of the Src family (SFK). We thus also evaluated the effects of dasatinib, a potent SFK inhibitor. We have shown previously that 100 nM dasatinib treatment completely blocked SFK activity in a few minutes (42) and we thus used this dose in all experiments presented in this study.

The efficiency of all inhibitors in blocking PAK kinase activity was assessed through the analysis of the extent of phosphorylation at the autophosphorylation site, which is present in both PAK1 (Ser144) and PAK2 (Ser141). Fig. 11A,B shows changes in phosphorylation of Ser144/141 in cells treated for 30 min with different IPA-3 concentrations. An example of the detected chemiluminiscence signal is shown in Fig. 11A. In HEK293T cells, the effect of IPA-3 was evaluated separately for each PAK1 band defined in Fig. 1. We noted considerable difference in IPA-3-induced dephosphorylation between cells treated in suspension and cells treated in adhered monolayer (Fig. 11B). The effect of IPA-3 on Ser144/141 phosphorylation was higher in cells treated in suspension (left). In addition, the sensitivity of the individual PAK1 bands to IPA-3 decreased with increasing apparent MW. The control inhibitor, PIR3.5, did not induce PAK dephosphorylation at Ser144/141 (Fig. S6A). In HeLa cells, PAK1 was barely detectable and it was difficult to quantify any change in PAK1 phosphorylation. Nevertheless, the trend was similar as in HEK293T cells (Fig. S6B). Quite surprisingly, the effect of SFK inhibition by dasatinib on Ser144/141 phosphorylation was only moderate and it was comparable for all PAK isoforms and conditions (Fig. 11C). We have also noted that the decrease in Ser144 phosphorylation was associated with an increase in PAK1 phosphorylation at both T212 and Ser20 in HEK293T cells treated with IPA-3 (Fig. S7).

**Fig. 11:**
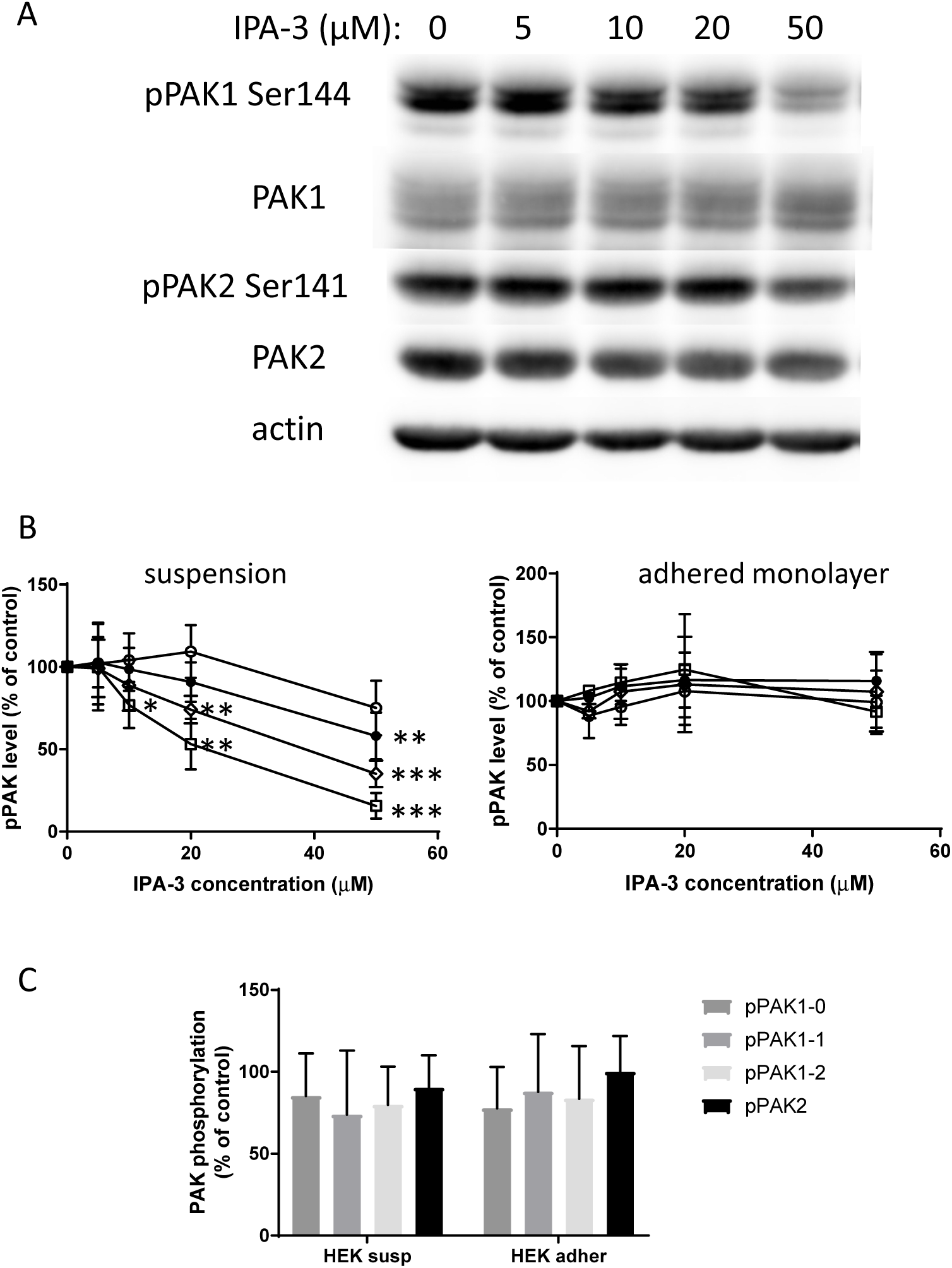
IPA-3 and dasatinib effect on PAK Ser144/Ser141 phosphorylation. HEK293T were treated for 30 min with different concentrations of IPA-3 or with 100 nM dasatinib. A: Example of western-blot analysis of Ser144/141 phosphorylation in IPA-3-treated samples. B: Summary and statistical evaluation of repeated experiments for IPA-3-treated HEK293T cells. The individual phospho-PAK bands are defined in Figure 1. Open symbols: circles - pPAK1-0, diamonds - pPAK1-1, squares – pPAK1-2. Closed symbols: pPAK2. Left: HEK293T cells treated in suspension, right: treated as adhered monolayer. C: Summary of results obtained for dasatinib-treated HEK293T cells, in suspension or in adhered monolayer as indicated. Differences between treated samples and controls were evaluated by paired t-test. * p < 0.05, ** p < 0.01, *** p < 0.001

To estimate possible cell damage due to the generated oxidative stress, we analyzed changes in the cell metabolism induced by the inhibitors used. Fig. 12 shows the effect of 1 h treatment with IPA-3 or PIR3.5 on the oxygen consumption rate (OCR) and on the rate of acidification outside the cells (ECAR), which reflects the glycolytic rate. The control compound PIR3.5 slightly reduced the respiration rate in HeLa cells, and a compensative increase in the glycolytic rate was observed in this cell line. In contrast, IPA-3 caused significant decrease of the respiration rate as well as of the glycolysis rate, in both cell lines. Similar results were obtained after 3h incubation with the inhibitors. On the other hand, no decrease in the cell metabolic rates occurred after 1 h dasatinib treatment (Fig. S8).

**Fig. 12:**
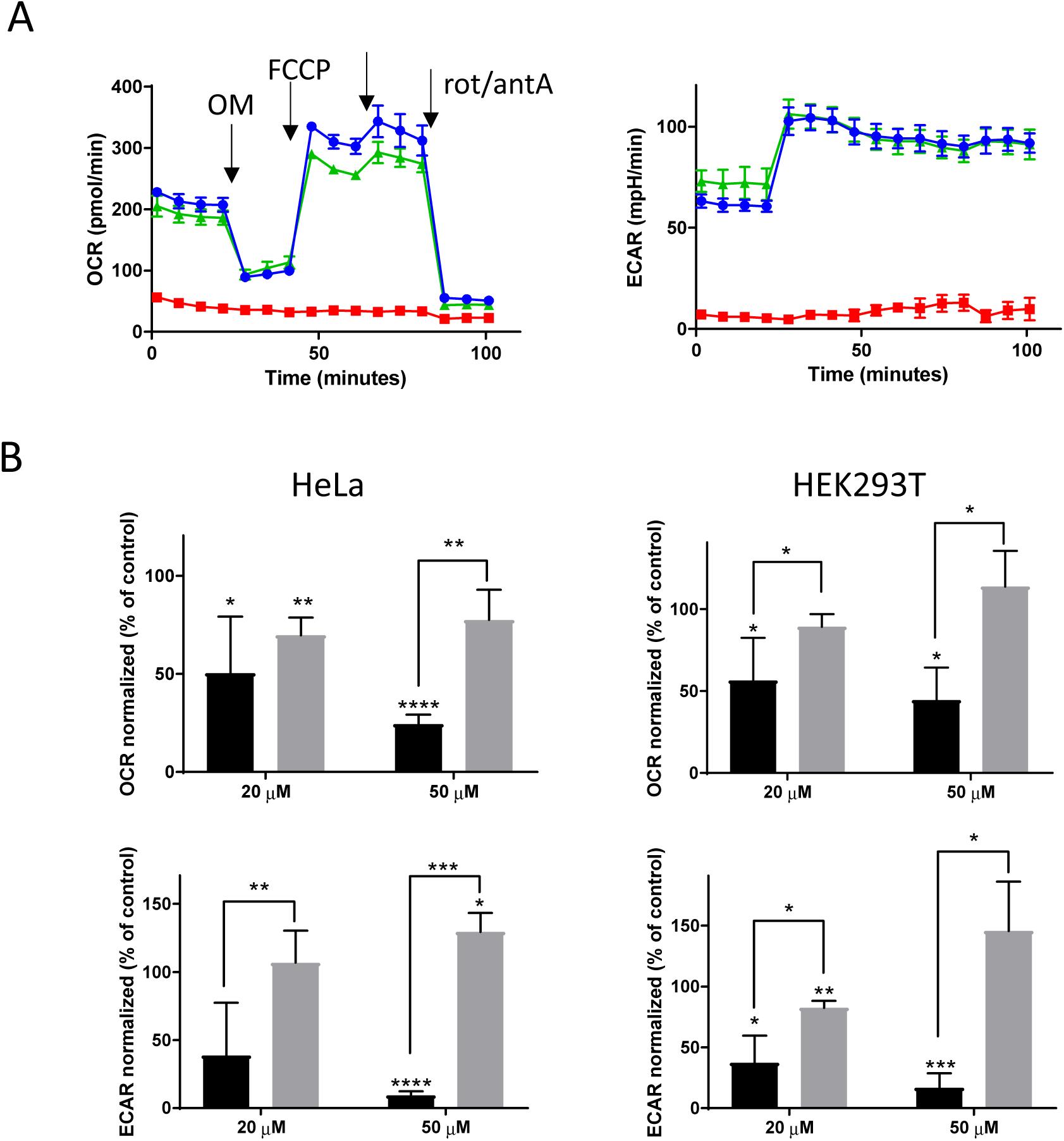
IPA-3 and PIR3.5 effect on the cell metabolism. The oxygen consumption rate (OCR) and the extracellular acidification rate (ECAR) were measured after 1 h cell pretreatment with IPA-3 or PIR3.5. A: Representative records for HeLa cells pretreated with 50 µM inhibitors. Injections of oligomycin (OM), FCCP, and rotenone/antimycin (rot/antA) are indicated by the arrows. Blue: control, green: PIR3.5, red: IPA-3. B: Summary values for the normalized basal metabolic rates before OM treatment. Upper plots: OCR, lower: ECAR. Black: IPA-3, grey: PIR3.5. Means and s.d. from 4 independent experiments. Left column: HeLa cells, right: HEK293T cells. The differences between normalized values in inhibitor-treated and control samples were evaluated using paired t-test. * p < 0.05, ** p < 0.01, *** p < 0.001

### Monitoring of cell-surface interactions

PAK group I are known to be involved in regulation of cell adhesion and migration. PAK1, as a downstream effector of SFK, promotes formation of membrane protrusions and releases the cytoskeletal tension, which is associated with a high stability of cell-surface adhesion points. The role of PAK2 in adhesion processes is much less explored, although differences between PAK1 and PAK2 have already been reported. Electrical Cell-Substrate Impedance Sensing (ECIS) is a powerful technique for real-time monitoring of cell interaction with a planar surface and we thus used this non-invasive method to study the cell response to inhibition of PAK or SFK activity. In the ECIS assay, gold microelectrodes embedded in the bottom of the testing plate enable to monitor cell binding to the well bottom through changes of the electrical impedance, which is further decomposed into resistance and capacitance. Typical examples of ECIS records are given in Fig. 13. The measurement was performed in two different settings: The cells were either pretreated with the inhibitors for 30 min before seeding into the ECIS plate (setting 1) or treated after the cell attachment, during signal recording (setting 2). In the latter case, the time of inhibitor addition is marked with an arrow. The largest changes occurred after dasatinib treatment and were indicative of decreased cell spreading. Indeed, microscopic analysis showed significant reduction of the cell area following dasatinib treatment in both cell lines (Fig. S9). On the other hand, the effects of PIR3.5 were only mild: we usually observed faster signal growth in HEK293 cells pretreated with PIR3.5 (setting 1) compared to the controls treated with solvent only. No reproducible change was detected in the setting 2, and no marked change was detected in HeLa cells treated with PIR3.5 under any condition (Fig. S10).

**Fig. 13:**
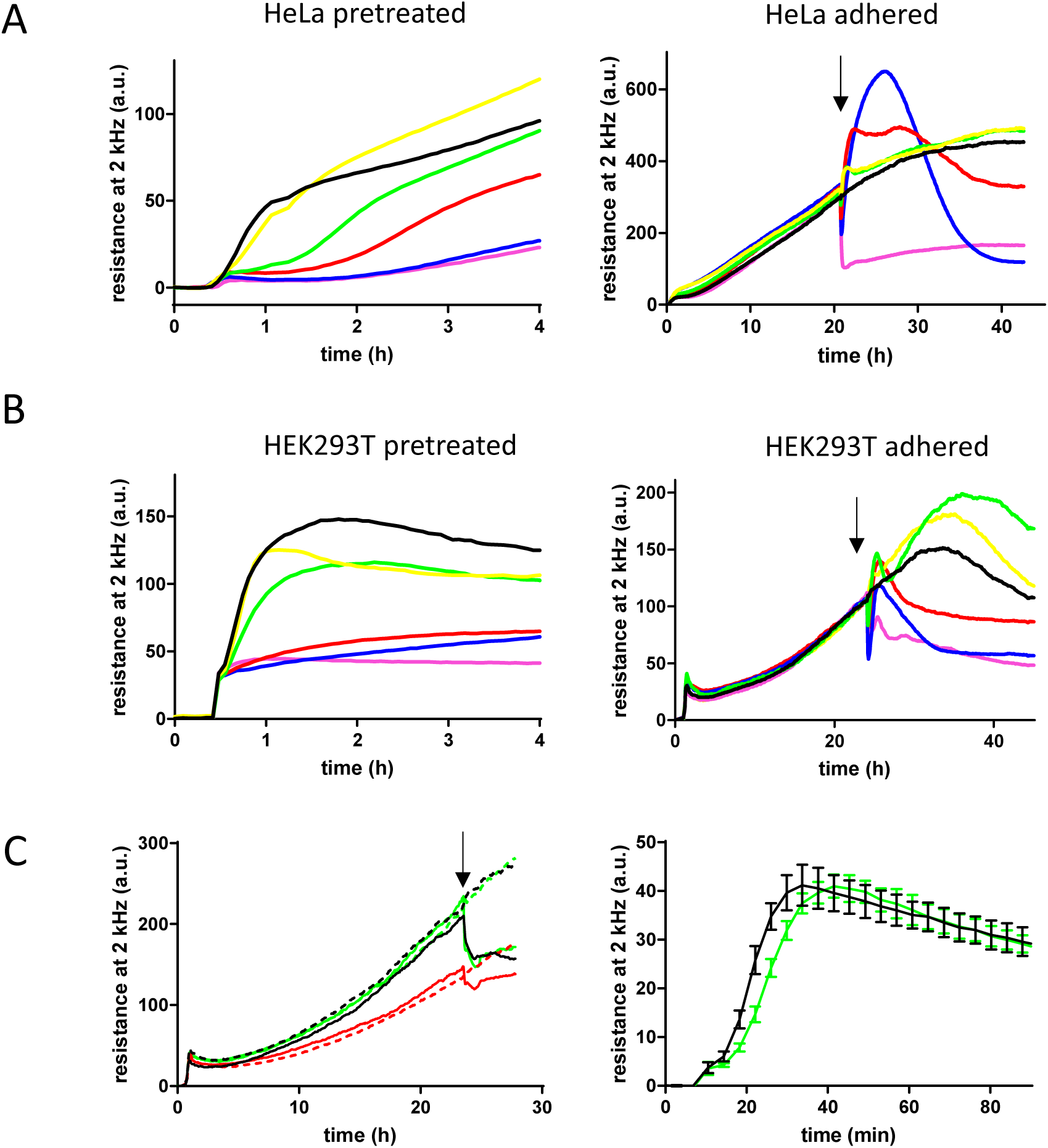
Representative records from cell attachment monitoring by ECIS. HeLa cells (A) or HEK293T cells (B) were preatreated for 30 min with inhibitors before seeding to ECIS wells (left) or treated during measurement (right). The arrows mark the time of inhibitor addition. Color legend: controls: black, IPA-3 at 5 – 10 – 20 – 50 µM: yellow – green – red – blue, dasatinib 100 nM: magenta. C: Attachment of HEK293T cells transfected with siRNA PAK1 or PAK2. Black: non-targeting control, red: siRNA PAK1, green: siRNA PAK2. In the left image, 100 nM dasatinib was added to the wells, which are represented by solid lines, at the time point indicated by the arrow.

The impact of IPA-3 on the microimpedance signal was cell-type dependent and time dependent. In HeLa cells, pretreatment with IPA-3 (setting 1, left) induced a progressive delay in cell attachment (Fig. 13A). At the highest dose (50 µM), the effect of IPA-3 approached that of dasatinib. In the setting 2 (right), IPA-3 up from 20 µM induced a distinct increase of the signal followed by a decrease, forming a large peak. More complex course occurred in HEK293T cells, which have higher PAK1 content. Cell pretreatment with low-dose IPA-3 did not prevent rapid cell attachment, but reduced the subsequent cell spreading. At 50 µM concentration, the effect of IPA-3 pretreatment was similar as in HeLa cells, close to that of dasatinib (Fig. 13B, left). In the setting 2 (Fig. 13B, right), an immediate signal drop occurred after IPA-3 addition, followed by one or two slower peaks. Although a small drop was usually also detectable in HeLa cells, its amplitude was significantly smaller (mean reduction by 60 % in HEK293T versus 18 % in HeLa cells after 20 µM IPA-3 addition, p = 0.008 from unpaired parametric t-test with Welch‘s correction, p = 0.0357 from Mann-Whitney test).

Fig. 13C illustrates the evolution of the microimpedance signal in HEK293T cells treated with siRNA to reduce PAK1 or PAK2 expression. The adhesion experiments were performed 24 or 48 h after siRNA transfection. The same cell number was seeded for all samples as checked by fluorescent staining of quadruplet sample aliquots. In agreement with the expected reduction of cell spreading, the amplitude of ECIS signal was lower for cells treated with PAK1 siRNA. Also, the fast drop of the signal after dasatinib addition was smaller in cells transfected with PAK1 siRNA: the mean decrease from 7 experiments was by 32 % in the non-targeting control and by 22% in PAK1 siRNA samples, the reached p-value was 0.05 from paired two-tail t-test. PAK2 silencing had no effect on the amplitude of the signal, but we noted a small delay during the first phase of the cell attachment (Fig. 13C, right). This effect was less marked in comparison with changes induced by siRNA PAK1, maybe due to lower efficiency in reducing the activity of PAK2. Nevertheless, similar effect was present in 5 of 6 experiments performed 48 h after the cell transfection with siRNA PAK2.

PAK1 splicing was reported to be mediated by the splicing factor U2AF65, the activity of this factor being modulated by the lysyl-hydroxylase JMJD6. JMJD6 silencing in melanoma cells resulted in an increased transcript level of the PAK1Δ15 compared to the full-length form (39). We thus tested the effect of JMJD6 silencing by siRNA in HEK293T cells on PAK expression at the protein level and on its phosphorylation, as well as on the cell adhesion. Representative results from repeated experiments are shown in Figure S11. The pattern of PAK1 bands was modified, the highest one being markedly reduced. However, the overall PAK1 expression level did not dramatically change and no additional band attributable to PAK1Δ15 was detected as a result of JMJD6 silencing. On the other hand, a loss of Ser144/141 phosphorylation was evident on PAK1 and, to a lesser extent, on PAK2. Accordingly, largely lower amplitude of the ECIS signal was observed in cells treated with JMJD6 siRNA.

## Discussion

PAK group I were discovered as effectors of small GTPases of the Rho family, which are the principal regulators of processes associated with cytoskeleton dynamics, such as cell adhesion and migration. In particular, the founding member of the PAK family, PAK1, has been analyzed in detail in this context. To date, PAK1 has well-established roles in the formation of plasma membrane protrusions (43–45), actin stress fiber dissolution and focal adhesion reorganization (46). PAK1 kinase activity appears necessary for disassembly of focal adhesions and of actin stress fibers, whereas membrane ruffling and lamellipodia formation are kinase independent (47).

Theoretically, multiple protein isoforms can be formed from the PAK1 gene. Both Swissprot and PubMed annotate two splicing isoforms, which give raise to the full-length PAK1 (553 AA) and to the variant lacking the exon 15, which is denoted as PAK1Δ15 (545 AA). In the majority of studies, these variants are not distinguished, despite possible differences in their functions. The ratio between PAK1 and PAK1Δ15 transcripts was reported to be regulated by the demethylase/hydroxylase JMJD6 in melanoma (39). In the same study, increased JMJD6 expression was associated with more aggressive disease and with worse survival in melanoma patients. JMJD6 was found to increase the PAK1/PAK1Δ15 ratio, to enhance MAPK signaling and to promote proliferation and invasion of melanoma cells.

PAK2 is known to be involved in the apoptosis (24, 48, 49). However, only a limited number of studies focused on specific functions of PAK2 in cell adhesion. The roles of PAK1 and PAK2 were analyzed in detail using siRNAs in a lung carcinoma model (8). Both PAK1 and PAK2 were required for heregulin-induced cell invasiveness, but many differences were observed in the mechanisms of their functions. PAK1 mediated cofilin dephosphorylation, and had stronger effect on lamellipodia formation. PAK2 was involved in the formation of new focal adhesions after heregulin stimulation and suppressed the activity of RhoA. PAK1 and PAK2 also had opposed effects on the myosin light chain phosphorylation.

In the present work, we aimed to characterize PAK1-full, PAK1Δ15 and PAK2 using several complementary approaches. We prepared plasmids for exogenous expression of all these kinases and compared their intracellular localization (Figs 8 and 10). Furthermore, we used the eGFP- and mCherry-tagged isoforms to analyze their mutual interactions using co-immunoprecipitation (Fig. 6). Functional differences between PAK1 and PAK2 were assessed using siRNA-mediated reduction of protein expression and by comparison of two cell lines, HEK293T and HeLa, the latter having much lower PAK1 expression than the former (Fig. 9).

Antibodies against PAK1 detected at least three different bands in western-blots (Figs 1, 2 and 4), that could correspond to splicing variants, but also to different activation stages. The electrophoretic mobility of recombinant PAK1 was described to change after *in vitro* activation by Cdc42 (10, 46). The existence of several PAK1 bands is generally believed to be due to multiple phosphorylation events occurring during the kinase activation. PAK1 has at least seven autophosphorylation sites (Ser21, Ser57, Ser144, Ser149, Ser199, Ser204 and Tyr423), and thirteen phosphorylated residues were found in a mass-spectrometry screen (50). However, although equivalent phosphorylation sites are for the most present on PAK2, this protein is detected in a single dominant band. To check the association between PAK1 phosphorylation and the band position on western-blots, we used alkaline phosphatase (AP) treatment (Fig. 5). The extent of phosphorylation at Ser144 and at Thr212 was markedly reduced by overnight treatment with 2250 U of AP. Although the majority of PAK bands were actually slightly shifted to a lower apparent MW, they remained distinctly separated. This result suggests that the hyperphosphorylation is not the main cause of the band multiplicity. Instead, other posttranslational modification could occur during PAK1 activation, possibly similar to PAK2 myristoylation upon caspase-mediated cleavage (40). We have also noted that the affinity of some antibodies to PAK1 seems to be affected by phosphorylation (e.g. ab223849, Fig. 5). Due to the complex band pattern in cells with exogenous PAK1 expression, it is not clear if the naturally occurring bands are derived from PAK1-full or PAK1Δ15. A mix of both may be present, the band position being further influenced by protein activation and phosphorylation. As the bands at higher MW were largely reduced by JMJD6 silencing (Figure S11), they could originate from PAK1-full. On the other hand, the position of the lowest band seems to correspond to that of the exogenous PAK1Δ15 (Figs 4 and 5).

The intracellular localization of the eGFP-tagged PAK1-full is in agreement with the current knowledge about PAK1 function, especially with the stimulation of membrane protrusions at the leading edge of migrating cells (Fig. 8B). On the other hand, the localization of PAK1Δ15 was similar to that of PAK2: both these kinases were enriched in focal adhesions (FA) (Figs 8A, 8B and 10A). With regard to the sequence comparison shown in Fig. 3, targeting to FA could be related to the C-terminal sequence. Interestingly, PAK1 was found in FA in some of the previous works (18, 46). Although the splicing variant was not explicitly defined, the PAK1 form used in these experiments was reported to have 544 AA, and was thus probably derived from PAK1Δ15. The kinase was released from FA upon Akt-mediated phosphorylation at Ser21 (18). PAK2 presence in FA detected in our experiments (Figs 8 and 10) corresponds to its reported role during assembly of new adhesion points (8).

Co-immunoprecipitation experiments confirmed that both PAK1 and PAK2 form homodimers (Fig. 6). In addition, we have shown that PAK1-full, PAK1Δ15 and PAK2 can mutually interact in all possible combinations. It is thus likely that they also mutually affect their function. In line with this hypothesis, we noted that in cells co-transfected with PAK2/PAK1Δ15 and PAK1-full, only few cells expressed high levels of both kinases simultaneously. The localization of PAK2/PAK1Δ15 (green) in FA was more apparent in the absence of PAK1-full (red) overexpression (Fig. 8B). Also, PAK2 presence in FA was clearly detected in HeLa cells, but not in HEK293T cells, which express endogenous PAK1. Moreover, negative regulation of PAK1 by PAK2 have been suggested in epithelial cells, where PAK2 knock-down increased PAK1 phosphorylation (51).

In the co-immunoprecipitation experiments, we also observed truncation of PAK1Δ15, as well as of PAK2, during interaction with PAK1-full (Fig. 6A,B; Table 2). The fragment size was similar to that produced by caspase-mediated cleavage of PAK2. However, the process was probably caspase-independent, as it was unaffected by the inhibitor Q-VD-OPh (Fig. S3), which completely blocks caspase activity at the concentration used (52). Also, PAK1Δ15 sequence does not contain the consensus caspase cleavage site. We cannot exclude the possibility that the truncation was an artifact due to the fluorescence labeling of the PAK molecules, which could result in a steric hinderance altering the dimer conformation.

However, virtually identical PAK2 truncation was observed with the inverse labeling. Also, PAK2 was cleaved upon interaction with PAK1-full (Fig. 6A), but not with PAK1Δ15 (Fig. 6C), despite the identical N-terminal part of these PAK1 isoforms (Fig. 3). In any case, if the heterodimer-induced cleavage occurred *in vivo*, the removal of the N-terminal part of PAK2/PAK1Δ15 could have similar consequence as the apoptotic PAK2 cleavage, i.e. constitutive activation of the kinase due to the absence of AID. On the other hand, PAK1Δ15 lacks the myristoylation sequence which could redirect the kinase to cell membranes, as it is the case for caspase-cleaved PAK2 (40).

Cell response to an acute PAK inhibition was studied using the relatively specific inhibitor IPA-3 (53), which binds to the closed conformation of the protein and prevents kinase activation. Accordingly, the effect of IPA-3 on Ser144/141 phosphorylation was larger for PAK1 bands at lower MW (Fig. 11B), which should correspond to less activated forms (46). Nevertheless, specific effects of IPA-3 on cell adhesion (Fig. 13) and on metabolic rates (Fig. 12) occurred at lower concentrations than those required for noticeable PAK dephosphorylation at Ser144/141. PAK activation is a multistage process and Ser144/141 phosphorylation stabilizes the final open conformation (15). However, PAK1 acts also as a scaffold, and this function may be kinase-independent. Indeed, induction of membrane ruffling by PAK1 overexpression was reported to be largely independent of PAK1 kinase activity (3). IPA-3 treatment also induced an increase in phosphorylation at Ser20 and this effect was already apparent at the lowest IPA-3 doses used (Fig. S7). As it was expected, the control compound PIR3.5 did not induce changes in PAK phosphorylation. The observed effects are thus mostly attributable to PAK inhibition and not to an increased oxidative stress. The set of our experiments involving IPA-3 and siRNA (Fig. 13) indicates, that PAK1 regulates cell spreading, presumably by promoting actin remodelling and formation of membrane protrusions. On the other hand, PAK2 is required for FA assembly and PAK2 depletion/inhibition slows down the cell attachment to the surface. This is in agreement with results obtained in another cell type (8). The roles of PAK1 and PAK2 in cell adhesion are thus not redundant, although these kinases were reported to have virtually identical substrate specificity (54). It was suggested previously that PAK2 may functionally compensate for PAK1 in neovascularization and wound healing (55). On the other hand, mutually opposed functions of PAK1 and PAK2 were described for other processes, e.g. in paxillin targeting to extracellular vesicles (56). The possibility to document differences in functions between PAK1-full and PAK1Δ15 is limited as there is no mean of specific inhibition of these two forms, the exogenously produced proteins are mostly kinase-inactive, and they affect the function of the endogenous proteins through heterodimer formation (Fig. S4). However, our findings indicate that PAK1Δ15 is in some aspects similar to PAK2, rather than to PAK1-full. Measurement of the cell metabolism (Fig. 12) revealed that PAK inhibition markedly reduced both the respiration rate (OCR) and the glycolysis rate (ECAR). The oxidative stress produced by both IPA-3 and PIR3.5 might be partly responsible for decreased cell respiration. However, the cells treated with PIR3.5 were able to cover the energy deficit by increased glycolysis. In contrast, both respiration and glycolysis were inhibited in cells treated with IPA-3. Association between PAK activity and cell metabolism has already been noted in several previous works. PAK2 depletion resulted in a reduction of lactate production and of glucose consumption in human head and neck cancer cells (57). On the other hand, activated PAK1 was reported to inhibit the glycolysis in leukocytes by phosphorylation of the phosphoglycerate mutase, an enzyme of the glycolytic pathway (58). As the effect of IPA-3 in our experiments was similar in HEK293T and HeLa cells, which largely differ in PAK1 content (Fig. 9), PAK2 inhibition was probably responsible for the reduced glycolytic rate.

## Material and Methods

### Cell culture

HeLa and HEK293T cells were obtained as a gift and authenticated using analysis of short tandem repeats, the results were compared with ATCC database. The cells were cultured in the recommended medium (RPMI-1640 for HeLa, DMEM for HEK293T) with 10 % fetal calf serum, 100 U/ml penicillin and 100 µg/ml streptomycin at 37 °C in 5 % CO_2_ humidified atmosphere. Cells from resuscitated frozen aliquots were not passaged for more than 3 months.

### Inhibitors and antibodies

IPA-3 (#3622) and PIR3.5 (#4212) were purchased from Tocris Bioscience and dissolved in sterile dimethylsulfoxide (DMSO) to make 50 mM stock solutions. Working solutions were prepared by 10 fold dilution of the stock solution in 50 mM Tris, pH 8.0, immediately before use. The cell density was adjusted to 3×10^5^ cells/ml for all experiments involving IPA-3 and PIR3.5 treatment.

Dasatinib was obtained from Selleckchem (#S1021), 200 µM stock solution was made in sterile DMSO. The antibodies against PAK1/2 are specified in Table 1. Other antibodies were purchased from the following providers: JMJD6 (sc 28348) and GFP (sc-9996) from Santa Cruz, β-actin (A5441) from Sigma-Aldrich, PAK3 from Cell Signaling (#2609). Anti-RFP was from Santa Cruz (sc-390909) or from Chromotek (6G6).

### Plasmid preparation and cell transfection

DNA fragments coding for PAK1-full, PAK1Δ15, and PAK2 were amplified from cDNA library (Jurkat cells, Origene) by PCR, using merged primers containing appropriate restriction sites (PAK1, both isoforms, Fw: AAAAAAAAGCTTCATGTCAAATAACGGCCTAGACA; Pak1-full Rv: AAAAAAGGATCCCGCTGCAGCAATCAGTGGA; PAK1Δ15 Rv: AAAAAAGGATCCCGTGATTGTTCTTTGTTGCCTCC; PAK2 Fw: AAAAAACTCGAGCATGTCTGATAACGGAGAACTG, Rv: AAAAAAGGATCCCACGGTTACTCTTCATTGCTTCT). To obtain expression of full-length proteins with no tag, the stop codon TAA was introduced into the reverse primers for PCR amplification of both PAK1 isoforms. The fragments were inserted into vectors peGFP-N2 or pmCherry-N2 (originally Clontech) designed for expression of proteins tagged with eGFP and mCherry at the C-terminus by standard methods of molecular cloning. Resulting plasmids were amplified in E. coli and purified with the PureYield Plasmid Miniprep System (Promega). The plasmids were then transfected into HEK293T cells using jetPRIME transfection reagent (Polyplus Transfection) following the manufacturer’s instructions. ON-TARGETplus siRNA (Dharmacon) targeting PAK1 (#L-003521), PAK2 (#L-003597), or JMJD6 (#L-010363), in parallel with a non-targeting control (#D-001810), were transfected using the same reagent (jetPRIME). The concentration range for siRNA was 100-150 nM (final concentration during transfection). The cells were then cultivated for 24 to 48 h without changing medium and harvested for further analyses.

### Western-blotting

The cells, when in suspension, were pelleted by centrifugation, washed once with ice-cold HBS (HEPES – buffered saline; 20 mM HEPES, 150 mM NaCl, pH 7.1) and lysed for 10 min/4 °C in Pierce IP Lysis Buffer (#87787) with freshly added protease and phosphatase inhibitors. When cultured on a plate in adherent layer, the cells were washed directly in the plate and scrapped into the lysis buffer. The suspension was then transferred to a centrifugation tube and incubated for 10 min/4 °C. Cellular debris was removed by centrifugation (15.000 g/4 °C/15 min), the lysate was mixed 1:1 (v/v) with 2x Laemmli sample buffer and incubated for 5 min at 95 °C.

An equivalent of 20 µg of total protein was resolved on 7.5 % polyacrylamide gel (18×18 cm) and transferred to a nitrocellulose membrane. The membrane was blocked for 1 h in 3 % bovine serum albumin and incubated for 1 h with the primary antibody in PBS with 0.1 % Tween-20 (PBST), at the room temperature. Thereafter, it was washed in PBST six times and incubated with the corresponding HRP-conjugated secondary antibody for 1 h. The chemiluminiscence signal from Clarity Western ECL Substrate (BioRad, #170-5060) was detected and analyzed using G:BOX iChemi XT-4 (Syngene).

### Alkaline phosphatase (AP) treatment of cell lysates

The cells were lysed as described in the western-blotting section. After centrifugation, lysates were diluted five fold in AP buffer (5 mM TRIS HCl, 30 mM NaCl, 1.5 mM MgCl2, 0.2 % NP-40, 0.2 mM EDTA, 1 % glycerol; pH 8.0) and treated with ∼2250 U AP (for the protein yield from 1×10^6^ cells). The samples were incubated at 37 °C for 20 to 24 h. The reaction was stopped by addition of 2x Laemmli sample buffer followed by incubation at 95 °C for 5 min.

### Electrical cell-substrate impedance sensing (ECIS)

Impedance measurements were performed using the ECIS Zθ device (Applied Biophysics). The wells of a 8W10E+ plate were filled with 200 µl culture medium and the baseline was monitored for several hours before cell addition. In the setting 1, the medium was removed and 400 µl of cell suspension was added. In this setting, the cells were pretreated for 30 min with inhibitors. In parallel, aliquots of cell suspensions were used to check for equal cell numbers by fluorescent staining (CyQuant Cell Proliferation Assay Kit; Molecular Probes, #C7026). In the setting 2, the cell suspension (200 µl) was added to the wells, the cell attachment was monitored overnight, and the inhibitors were added after 20 to 24 h. One of the wells in each plate were left empty (medium only) and the signal was used as a baseline for the other wells. The instrument automatically decomposes the impedance signal into resistance and capacitance. The ECIS records were exported to Excel and processed using the GraphPad Prism software: the background was set to zero at a time point shortly before cell seeding, and the baseline (empty well) was subtracted. The signals shown in the graphs represent the averages from two identically treated plates, which were run in parallel.

### Confocal microscopy

Localization of PAK was analyzed by live cell imaging in cells transfected with plasmids for expression of proteins with fluorescent tags and by immunofluorescence in cells fixed with 2 % paraformaldehyde and permeabilized with 0.3 % Triton X-100. Interference reflection microscopy (IRM) was used to visualize cell-substrate contact points. The measurement was performed by means of FV-1000 confocal microscope (Olympus), using 405 nm laser beam and focusing to the glass surface.

### Immunoprecipitation

Immunoprecipitation using GFP- or RFP-Trap (Chromotek) was performed according to manufactureŕs instruction as described previously (59). Briefly, cells were harvested and washed with PBS, lysed in Lysis buffer (10 mM Tris/Cl pH7.5, 150 mM NaCl, 0.5 mM EDTA, 0.5 % NP-40, protease and phosphatase inhibitors) for 30 min/4 °C and centrifuged at 20.000 g/4 °C for 10 min. A small fraction of lysate was put aside for native and SDS-PAGE and the majority of lysate was mixed with GFP/RFP-nanobody-coated beads and rotated for 1 h/4 °C. Then the beads were extensively washed with diluting buffer (10 mM Tris/Cl pH 7.5, 150 mM NaCl, 0.5 mM EDTA), resuspended in Laemmli sample buffer (50 mM Tris pH 6.8, 2 % SDS, 100 mM DTT, 10 % glycerol), boiled at 95 °C for 10 min and centrifuged 20.000 g/4 °C for 10 min. Supernatant was stored at −20 °C until used for SDS-PAGE.

### Native and semi-native PAGE

Lysates obtained from the immunoprecipitation procedure were mixed with 2x native buffer (50 mM Tris pH 6.8, 10 mM DTT, 10 % glycerol) and without boiling subjected to 4-15 % gradient gel (BioRad) without SDS for native ELFO, or to 7.5 % polyacrylamide gel with 0.1 % SDS for semi-native ELFO. After blotting to PVDF membrane (BioRad), the blot was incubated with GFP primary antibody at a dilution 1:250 or 1:500 for native or semi-native samples. Anti-mouse HRP-conjugated secondary antibody from Thermo Scientific was used at concentrations 1:20.000 and 1:50.000, respectively. ECL Plus Western Blotting Detection System (GE Healthcare) was used for chemiluminiscence visualization and evaluation by G-box iChemi XT4 digital imaging device (Syngene Europe).

### Cell metabolism measurement

The oxygen consumption rate (OCR) and the extracellular acidification rate (ECAR) were measured using the Mito Stress Test kit (Agilent, #103010-100), following the manufactureŕs instructions. HeLa (16.000 cells/well) or HEK293T (24.000 cells/well) were seeded to the plates in 80 µl medium and incubated overnight. The plate was then washed with medium without fetal bovine serum, the cells were treated with inhibitors in dublets and incubated for 1 h at 37 °C, without CO_2_. The control cells were treated with the solvent (DMSO in Tris buffer) only. The plate was washed again with medium without serum and analyzed using the Seahorse XFp apparatus (Agilent). The final concentrations of the injected compounds were the following: oligomycin 1 µM, carbonyl cyanide-4 (trifluoromethoxy) phenylhydrazone (FCCP) 0.3 and 0.5 µM, rotenone/antimycin A 0.5 µM. After the measurement, the medium was aspirated and the cell number was assessed using CyQuant Cell Proliferation Assay Kit (Molecular Probes, #C7026). The records were normalized and analyzed using the Wave software (Agilent).

## Acknowledgment

The authors wish to thank M. Voráčová for expert technical assistance and to H. Čechová for cell line authentication. The work was supported by the Grant Agency of the Czech Republic (grant No 16-16169S) and by the Ministry of Health of the Czech Republic (project for conceptual development of the research organization No 00023736).

