## Supplemental Figures for "PAK1, PAK1Δ15, and PAK2: similarities, differences and mutual interactions"

Figure S1: **Sequence comparison of PAK isoforms**

Triple sequence alignment was performed using Clustal W software.

```
                          1-50
PAK1 full length          MSNNGLDIQDKPPAPPMRNTSTMIGAGSKDAGTLNHGSKPLPPNPEEKKK
PAK1 delta15              MSNNGLDIQDKPPAPPMRNTSTMIGAGSKDAGTLNHGSKPLPPNPEEKKK
PAK2                      MSDNG-ELEDKPPAPPVRMSSTIFSTGGKDPLSANHSLKPLPSVPEEKKP
                          51-100
PAK1 full length          KDRFYRSILPGDKTNKKKEKERPEISLPSDFEHTIHVGFDAVTGEFTGMP
PAK1 delta15              KDRFYRSILPGDKTNKKKEKERPEISLPSDFEHTIHVGFDAVTGEFTGMP
PAK2                      RHKIISIFSGTEKGSKKKEKERPEISPPSDFEHTIHVGFDAVTGEFTGMP
                          101-150
PAK1 full length          EQWARLLQTSNITKSEQKKNPQAVLDVLEFYNSKKTSNSQKYMSFTDKSA
PAK1 delta15              EQWARLLQTSNITKSEQKKNPQAVLDVLEFYNSKKTSNSQKYMSFTDKSA
PAK2                      EQWARLLQTSNITKLEQKKNPQAVLDVLKFYDSNT--VKQKYLSFTPPEK
                          151-199
PAK1 full length          EDYNS-SNALNVKAVSETPAVPPVSEDEDDDDDDATPPPVIAPRPEHTKS
PAK1 delta15              EDYNS-SNALNVKAVSETPAVPPVSEDEDDDDDDATPPPVIAPRPEHTKS
PAK2                      DGFPSGTPALNAKG-TEAPAV--VTEEEDD--DEETAPPVIAPRPDHTKS
                          200-249
PAK1 full length          VYTRSVIEPLPVTPTRDVATSPISPTENNTTPPDALTRNTEKQKKKPKMS
PAK1 delta15              VYTRSVIEPLPVTPTRDVATSPISPTENNTTPPDALTRNTEKQKKKPKMS
PAK2                      IYTRSVIDPVPA-PVGD-------------SHVDGAAKSLDKQKKKTKMT
                          250-299
PAK1 full length          DEEILEKLRSIVSVGDPKKKYTRFEKIGQGASGTVYTAMDVATGQEVAIK
PAK1 delta15              DEEILEKLRSIVSVGDPKKKYTRFEKIGQGASGTVYTAMDVATGQEVAIK
PAK2                      DEEIMEKLRTIVSIGDPKKKYTRYEKIGQGASGTVFTATDVALGQEVAIK
                          300-349
PAK1 full length          QMNLQQQPKKELIINEILVMRENKNPNIVNYLDSYLVGDELWVVMEYLAG
PAK1 delta15              QMNLQQQPKKELIINEILVMRENKNPNIVNYLDSYLVGDELWVVMEYLAG
PAK2                      QINLQKQPKKELIINEILVMKELKNPNIVNFLDSYLVGDELFVVMEYLAG
                          350-399
PAK1 full length          GSLTDVVTETCMDEGQIAAVCRECLQALEFLHSNQVIHRDIKSDNILLGM
PAK1 delta15              GSLTDVVTETCMDEGQIAAVCRECLQALEFLHSNQVIHRDIKSDNILLGM
PAK2                      GSLTDVVTETCMDEAQIAAVCRECLQALEFLHANQVIHRDIKSDNVLLGM
                          400-449
PAK1 full length          DGSVKLTDFGFCAQITPEQSKRSTMVGTPYWMAPEVVTRKAYGPKVDIWS
PAK1 delta15              DGSVKLTDFGFCAQITPEQSKRSTMVGTPYWMAPEVVTRKAYGPKVDIWS
PAK2                      EGSVKLTDFGFCAQITPEQSKRSTMVGTPYWMAPEVVTRKAYGPKVDIWS
                          450-499
PAK1 full length          LGIMAIEMIEGEPPYLNENPLRALYLIATNGTPELQNPEKLSAIFRDFLN
PAK1 delta15              LGIMAIEMIEGEPPYLNENPLRALYLIATNGTPELQNPEKLSAIFRDFLN
PAK2                      LGIMAIEMVEGEPPYLNENPLRALYLIATNGTPELQNPEKLSPIFRDFLN
                          500-548
PAK1 full length          RCLEMDVEKRGSAKELLQVRKLRFQ-VFSNFSMIAASIPEDCQAPLQPHS
PAK1 delta15              RCLEMDVEKRGSAKELLQHQFLKIAKPLSSLTPLIAAAK---------EA
PAK2                      RCLEMDVEKRGSAKELLQHPFLKLAKPLSSLTPLIMAAK---------EA
                          549-553
PAK1 full length          TDCCS
PAK1 delta15              TKNNH
PAK2                      MKSNR
```

Figure S2: **Detection of the truncated PAK2-eGFP in immunoprecipitates**
HEK293T cells were transfected with PAK2-eGFP in combination with PAK1-full-mCherry or with a control mCherry plasmid. The immunoprecipitates obtained from beads binding mCherry/RFP or GFP were probed by GFP antibody. The truncated PAK2 was not found in the absence of PAK1-full.

1. PAK2-eGFP + PAK1-full-mCherry, IP:RFP
2. PAK2-eGFP + PAK1-full-mCherry, IP:GFP
3. PAK2-eGFP + empty mCherry, IP:GFP

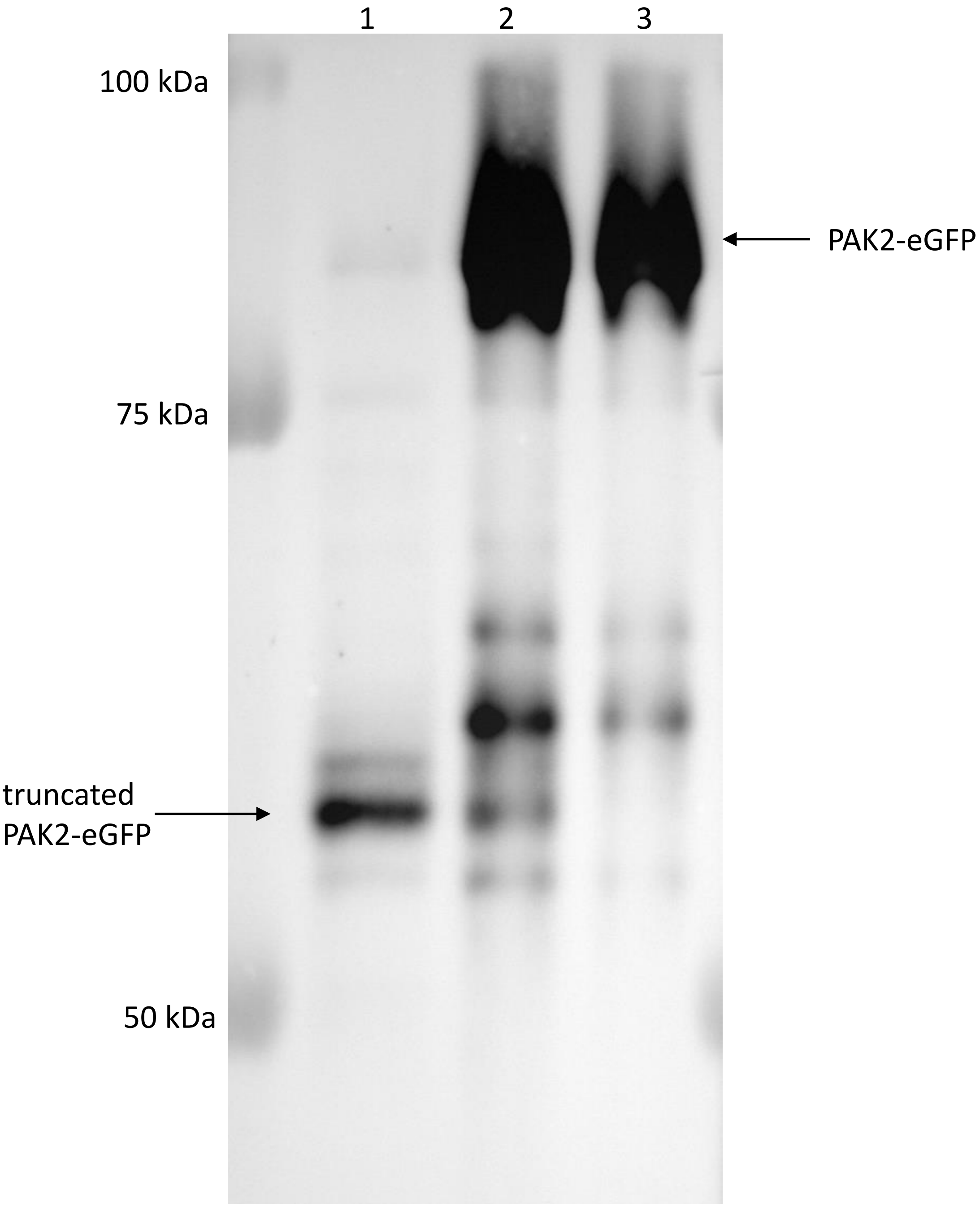

Figure S3: **Effect of caspase inhibition on PAK2-eGFP truncation**
HEK293T cells were transfected with the indicated plasmids in the absence or in the presence of 10 µM Q-VD-OPh (added 1 h prior to transfection and maintained until the cell harvest).

1 PAK2-eGFP + PAK1-mCherry
2 PAK2-eGFP + PAK1-mCherry + Q-VD-OPh
3 PAK2-eGFP + empty mCherry
4 PAK2-eGFP + empty mCherry + Q-VD-OPh

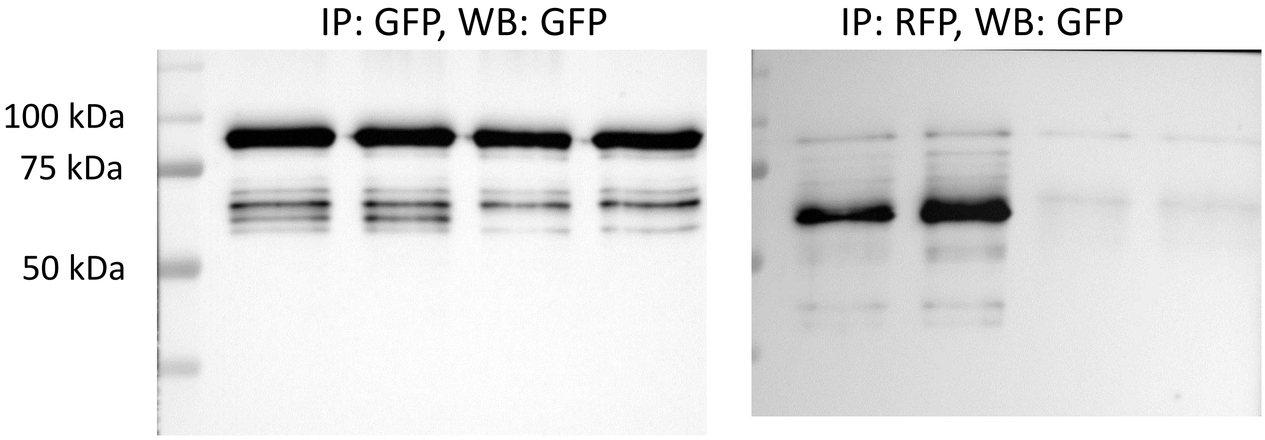

Figure S4: **Detection of the endogenous PAK1 in immunoprecipitates.**
HEK293T cells were transfected with the plasmid encoding PAK1-full-GFP. The exogenous PAK1 with its interaction partners was pulled down using GFP-trap beads. The precipitate was resolved using SDS electrophoresis and blotted to two membranes, which were incubated with antibodies against GFP (left) or PAK1 (ab223849, right). The position of MW markers is indicated on the left side. The blue arrow points to the expected position of the endogenous PAK1 (64-67 kDa). No band was detected at this position using GFP antibody.

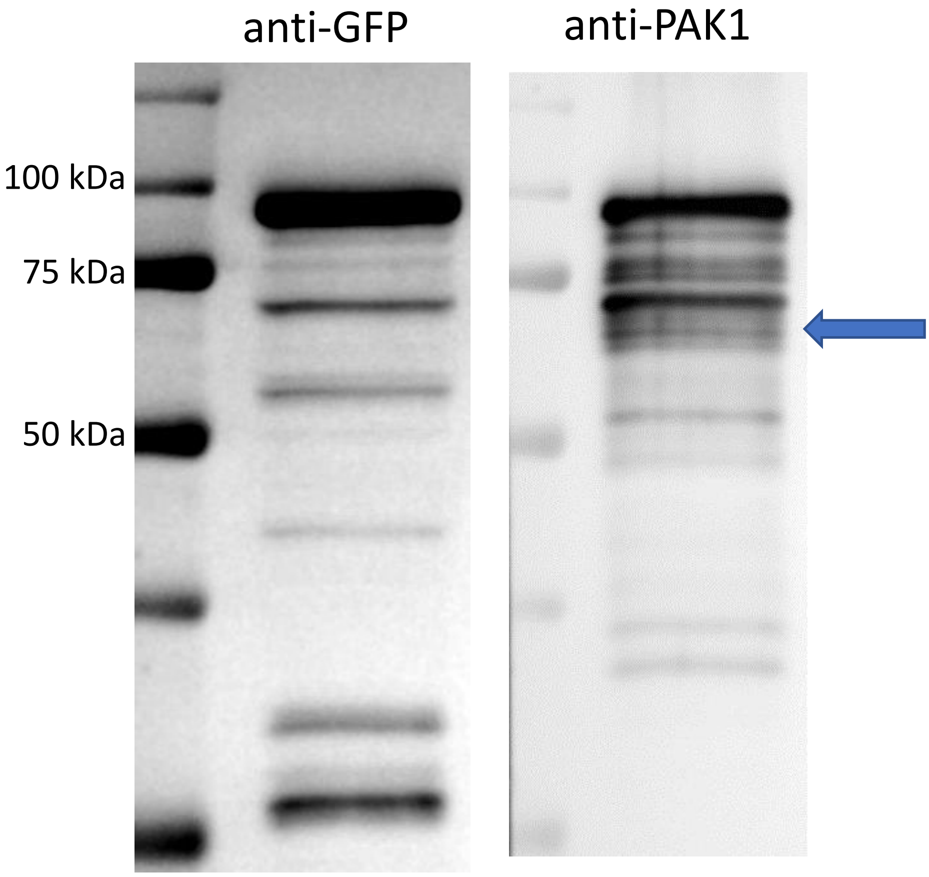

Figure S5: **Cell viability after 24 h treatment with IPA-3 and PIR3.5**
HeLa cells were harvested from plates at the end of ECIS measurement, 24 h after inhibitor addition (setting 2). Cell viability was measured by the standard propidium iodide (PI) exclusion assay using the flow-cytometer BD Fortessa. Means and s.d. from 5 to 9 experiments for each condition.

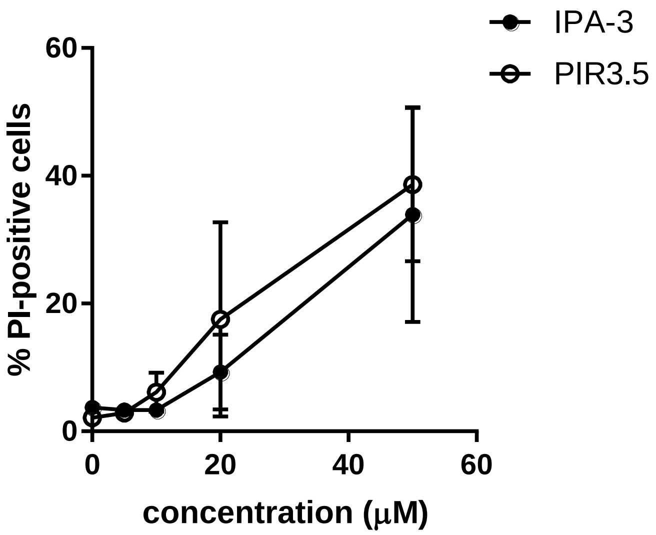

Figure S6: **IPA-3 or PIR3.5-induced changes in PAK Ser144/141 phosphorylation**
The cells were treated for 30 min with different concentrations of IPA-3 or PIR3.5. Means and s.d. from at least 3 repeated experiments for each condition. The individual phospho-PAK bands are defined in Figure 1. Open symbols: circles - pPAK1-0, diamonds - pPAK1-1, squares – pPAK1-2. Closed symbols: pPAK2.

A. HEK293T cells treated with PIR3.5 in suspension or as an adhered monolayer

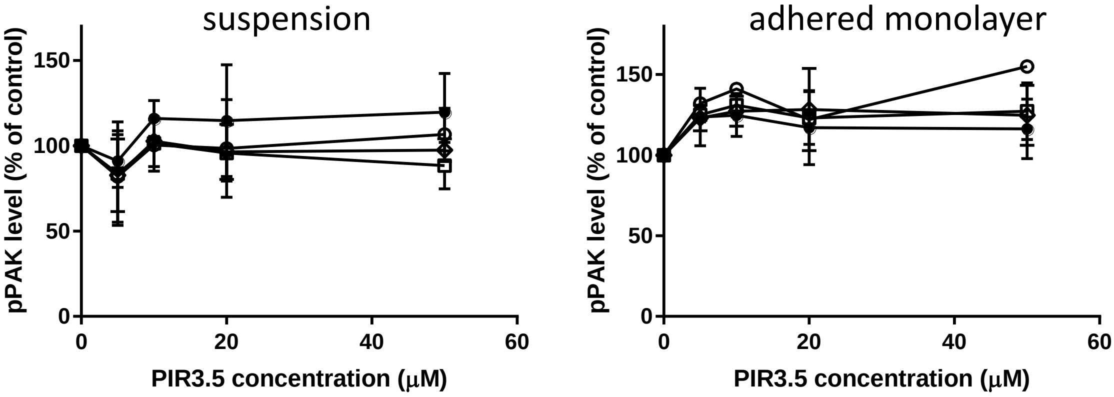


B. HeLa cells treated with IPA-3 (upper plots) or PIR3.5 (lower plots) in suspension or as an adhered monolayer

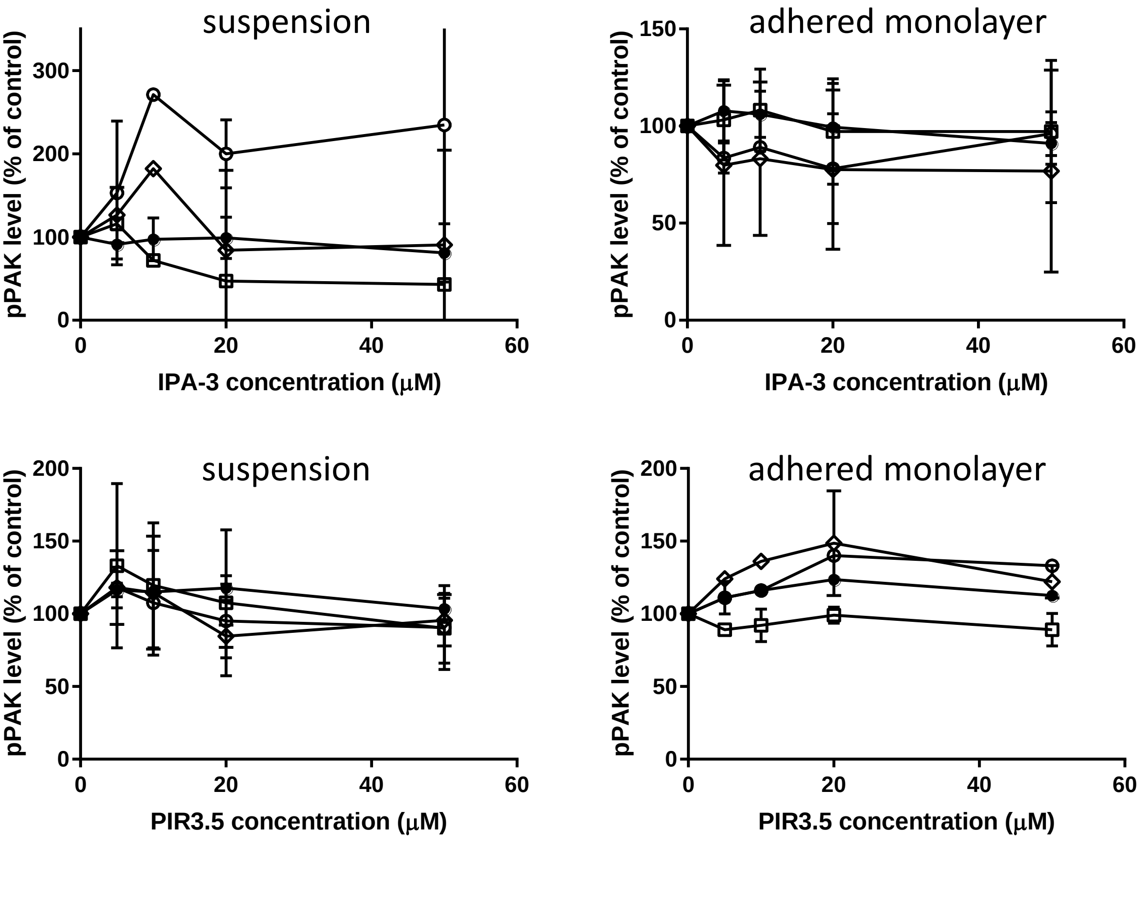

Figure S7: **IPA-3-induced changes in T212 and Ser20 phosphorylation**
HEK293T cells in suspension were treated for 30 min with IPA-3. The lysates were resolved by gel electrophoresis and the western-blot membranes were incubated with phospho-specific antibodies recognizing pT212 (ab75599) or pSer20 (ab51244) of PAK1. The figure shows examples of the signals (upper part) and the summary of the results obtained (means and ranges from 7 and 5 experiments for T212 and Ser20, respectively).

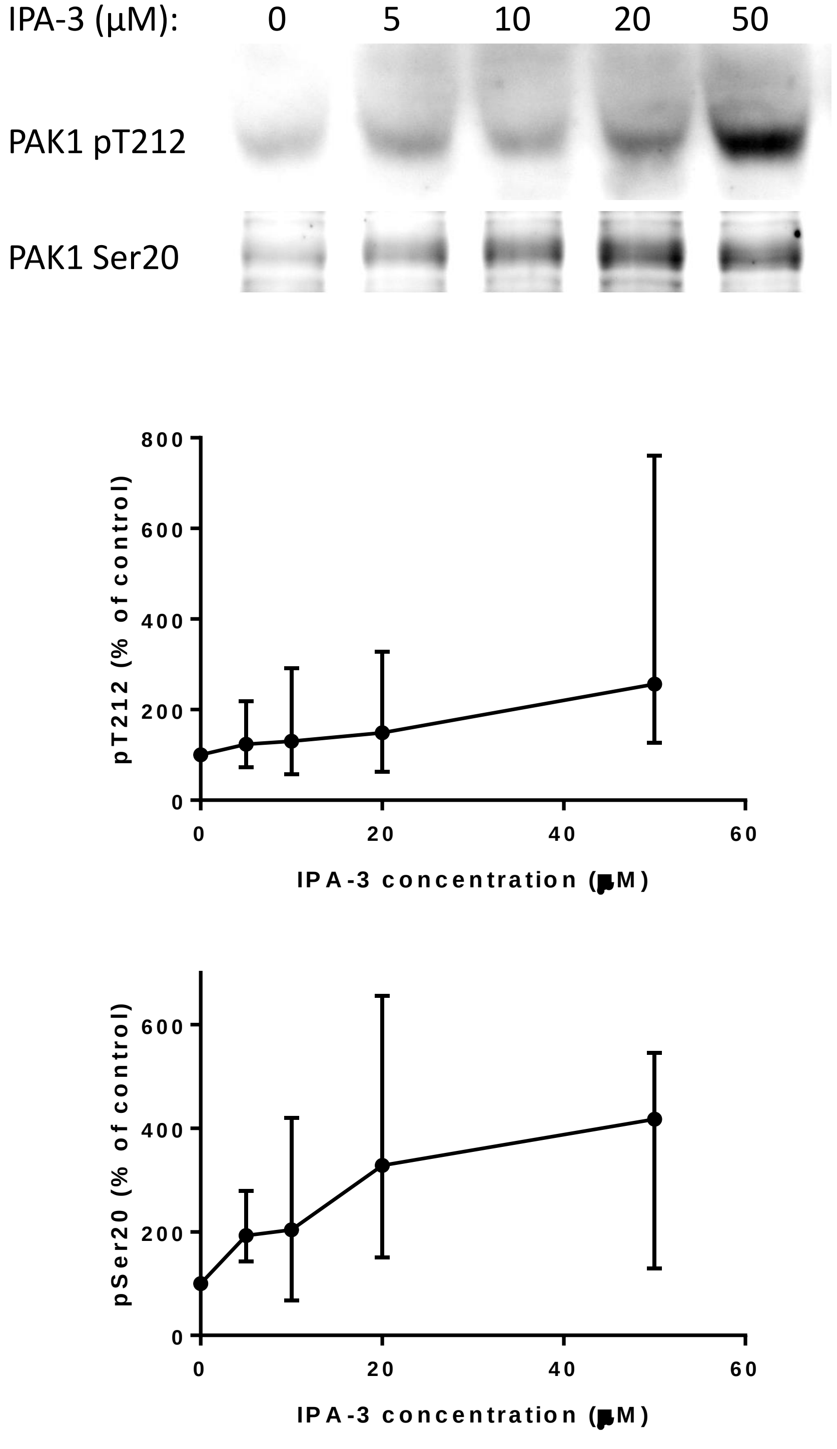

Figure S8: **Effect of 100 nM dasatinib on the cell metabolism**
The cells were treated for 1 h with 100 nM dasatinib and the oxygen consumption rate (OCR) and the extracellular acidification rate (ECAR) were measured using Seahorse XFp apparatus as described in Fig. 12. The graph shows means and s.d. of the basal metabolic rates from two experiments for each cell line.

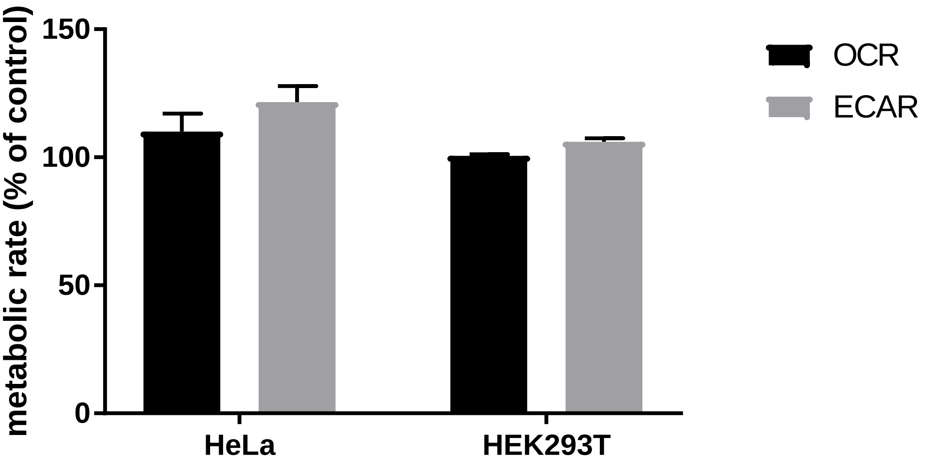

Figure S9: **Morphological changes induced by dasatinib**
Cells were treated for 90 min with 100 nM dasatinib and their morphology was examined using the Nanolive microscope (Agilent). A: HEK293T cells, representative examples of non-treated and treated cells. B: HeLa cells. The plot documents a significant decrease in the mean cell surface area after dasatinib treatment.

A. HEK293T

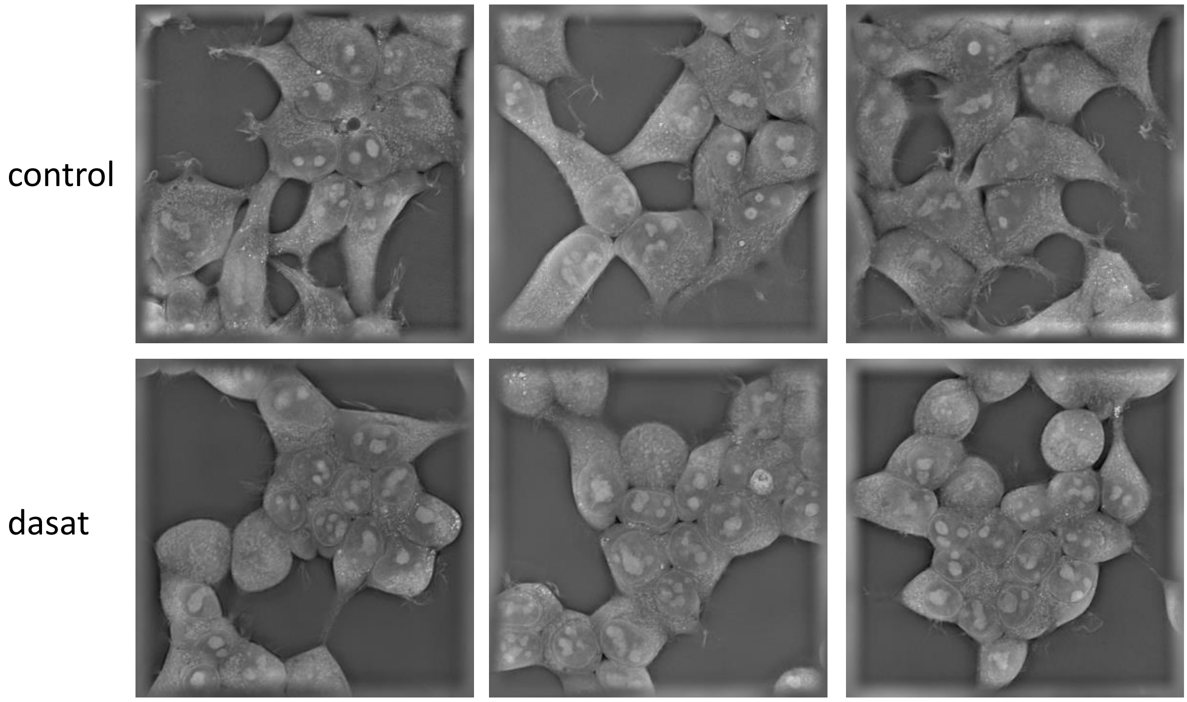


B. HeLa

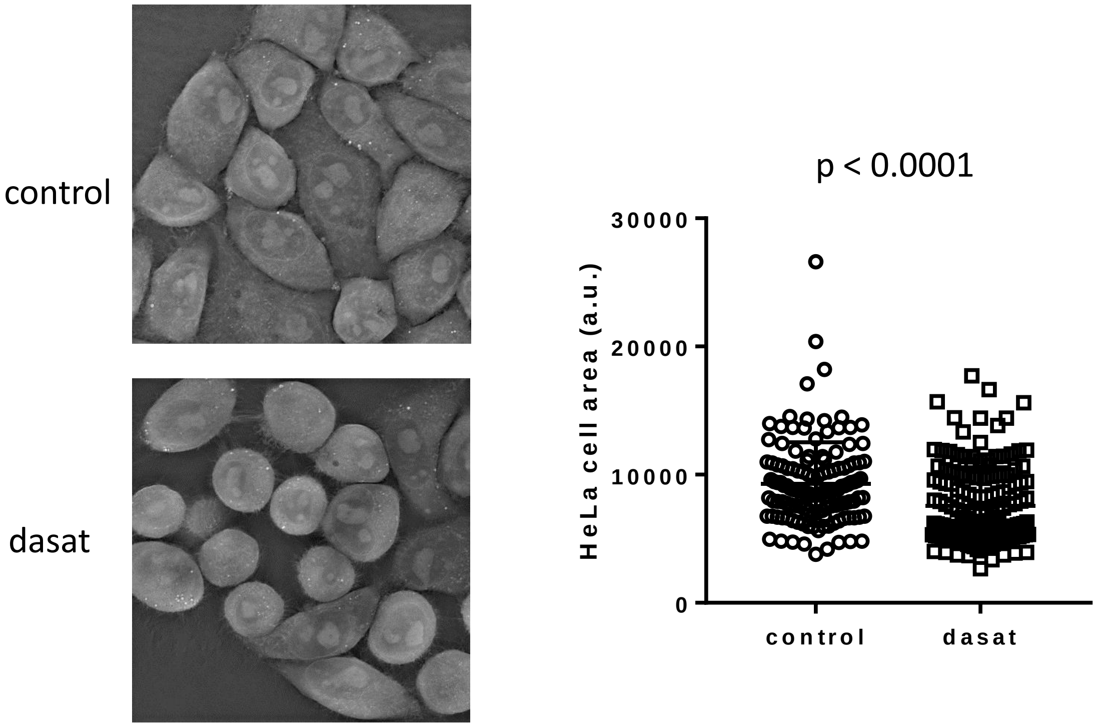

Figure S10: **Representative examples of ECIS records for PIR3.5 treatment**
HeLa cells (upper plots) or HEK293T cells (lower plots) were preatreated for 30 min with inhibitors before seeding to ECIS wells (left) or treated during measurement (right). The arrows mark the time of inhibitor addition. Color legend: controls: black, PIR3.5 at 5 – 10 – 20 – 50 µM: yellow – green – red – blue, dasatinib 100 nM: magenta.

HeLa

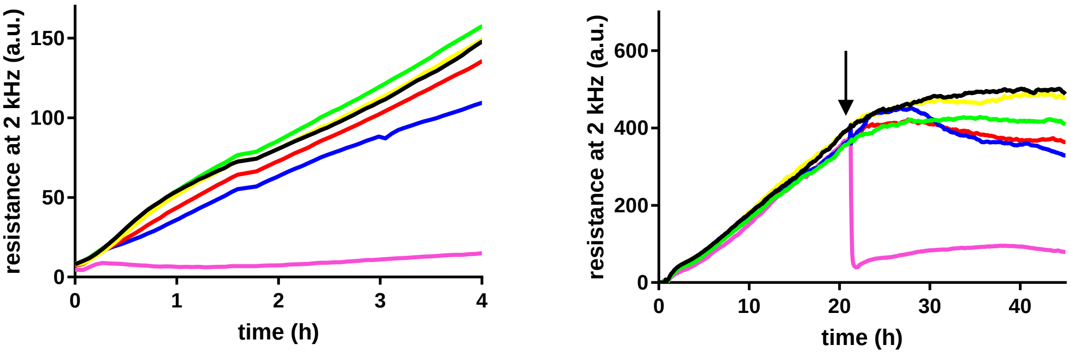


HEK293T

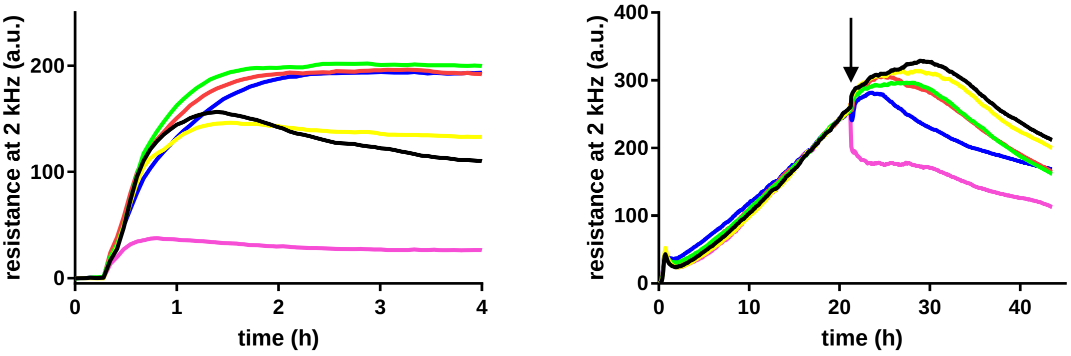

Figure S11: **Effect of JMJD6 silencing by siRNA**
HEK293T were transfected with siRNA JMJD6 and incubated for 48 or 72 h. JMJD6 and PAK expression was then analyzed by western-blotting and cell adhesion by ECIS (bottom). Green line: untransfected cells, red line: siRNA JMJD6.

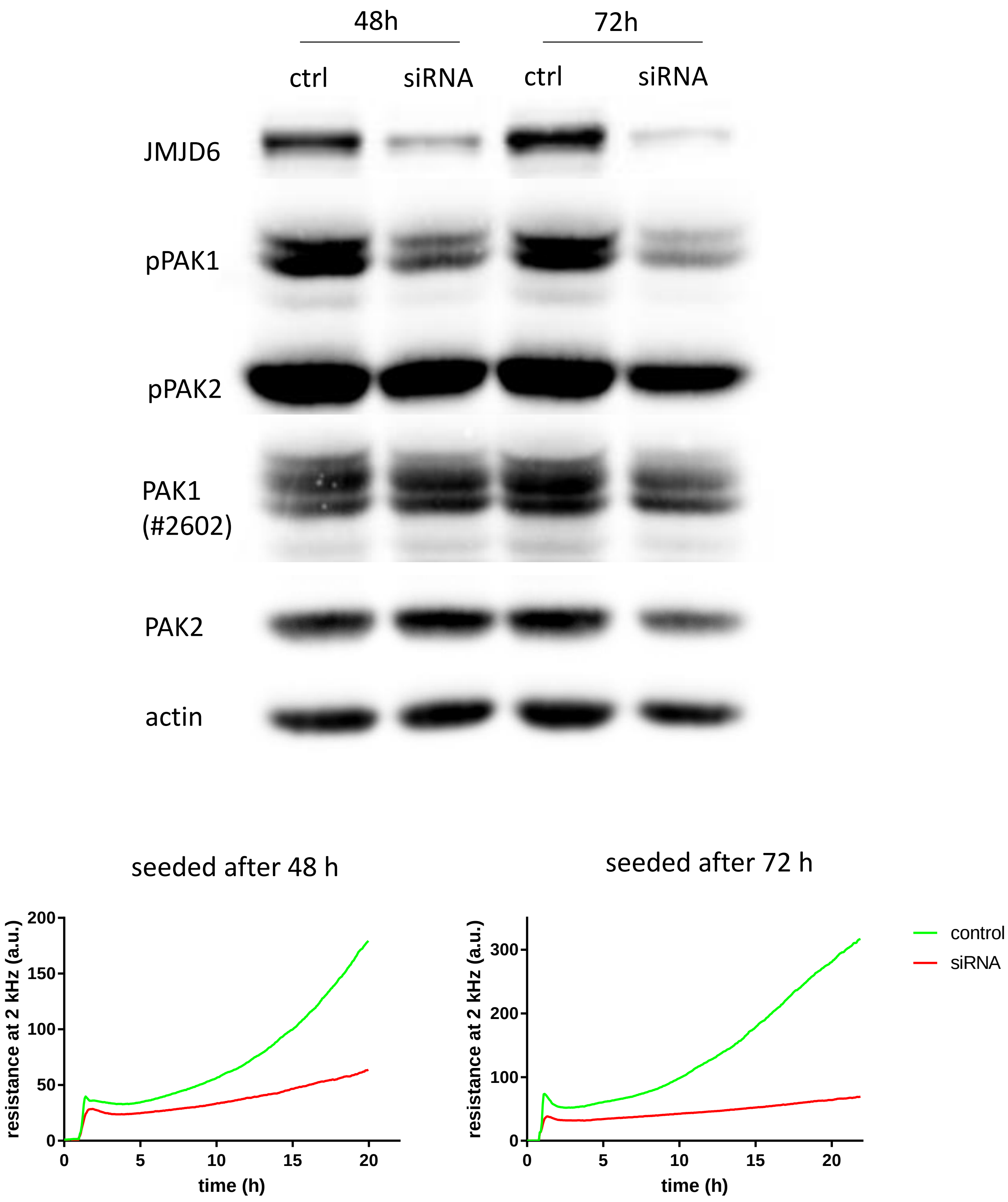
